# De novo design of cysteine proteases

**DOI:** 10.1101/2025.11.21.689808

**Authors:** Hojae Choi, Magnus S. Bauer, Brian Coventry, Preetham Venkatesh, Anqi Chen, Donghyo Kim, Asim K. Bera, Alex Kang, Hannah Nguyen, Shayan Sadre, Justin Decarreau, Emily Joyce, Bhanumathi Shankaran, Tuscan Rock Thompson, Gabriel Greenstein, Kieran Didi, Lars L. Schaaf, Jacob Merle Gershon, Alexander F. Shida, Gyu Rie Lee, Donald Hilvert, Samuel J. Pellock, David Baker

**Affiliations:** Department of Biochemistry, University of Washington, Seattle, WA 98195, USA; Institute for Protein Design, University of Washington, Seattle, WA 98195, USA; Graduate Program in Biological Physics, Structure and Design, University of Washington, Seattle, WA, USA; Howard Hughes Medical Institute, University of Washington, Seattle, WA 98195, USA; Molecular and Cellular Biology Program, University of Washington, Seattle, WA, USA; Molecular Biophysics and Integrated Bioimaging, Lawrence Berkeley National Laboratory, Berkeley, CA, USA; Department of Materials Science & Engineering, Stanford University, Stanford, CA 94305, USA; Department of Computer Science, University of Oxford, Parks Rd, Oxford OX1 3QD, UK; NVIDIA Corp., Santa Clara, USA; Cavendish Laboratory, Department of Physics, University of Cambridge, Cambridge CB3 0HE, UK; Department of Materials, Imperial College London, Exhibition Road, London SW7 2AZ, U.K; Department of Molecular Engineering, University of Washington, Seattle, WA, USA; Medical Scientist Training Program, University of Washington, Seattle, WA 98195, USA; Department of Biological Sciences, Korea Advanced Institute of Science and Technology, Daejeon, Republic of Korea, 34141; Laboratory of Organic Chemistry, ETH Zurich, Zurich, Switzerland

## Abstract

Despite advances in de novo enzyme design, success has been largely limited to low energy barrier model reactions. Amide bonds such as those linking amino acids along the peptide backbone are stable for hundreds of years in neutral aqueous solution because of the high energy barrier to hydrolysis^1^. Here we describe the de novo design of enzymes which utilize an activated cysteine nucleophile to hydrolyze the polypeptide backbone in a sequence-dependent manner, with a success rate of 13/69=19% and rate enhancements over the background reaction (*k*_cat_/*k*_uncat_) of up to 3 × 10^7^. The designed proteases have folds very different from proteases in nature (TM score < 0.50), and six crystal structures are very close to the design models (Cα RMSDs < 1.2 Å), highlighting the capacity for generalization and the accuracy of the design methodology. Experimental and computational analyses suggest that the remaining gap in activity to the most active native cysteine proteases arises from imperfections in active site preorganization and substrate positioning. The designed proteases efficiently cleave their targets in mammalian cells, opening the door to a wide range of synthetic biology applications.

## Introduction

Proteases – enzymes that cleave the amide bonds found in proteins – are widespread in nature and widely used in biotechnology^2^, yet their de novo design has remained elusive. This is likely due to the resilience of amides to hydrolysis: with half lives of hundreds of years^1^, they require substantial activation to be cleaved. The cysteine protease enzyme family overcomes this energy barrier by employing a hydrogen bonded aspartate-histidine-cysteine catalytic triad that activates the cysteine nucleophile, and an oxyanion hole that stabilizes multiple high energy intermediates and transition states along the multistep reaction pathway^3–6^. A peptide binding cleft with sequence specificity for the target substrate positions the scissile amide bond adjacent to the nucleophilic cysteine. While such catalytic machinery is well characterized in native enzymes, the complexity and polarity is a challenge for computational design, and has only recently been achieved for serine hydrolases that act on small non-polar ester substrates^7^ that are much more activated than the amides of proteins and easier to bind than polar protein substrates. Combined, the chemical stability of protein substrates and the complex architecture required to recognize and cleave them has made the de novo design of proteases a formidable design challenge that has remained unsolved for decades.

We reasoned that recent advances in ML-based protein design could construct the complex architecture necessary to cleave amide bonds and position the flexible peptide substrate. The generative AI model RFdiffusion2 (RFD2)^8,9^ enables the conditioning of diffusion trajectories on the positions of active site atoms without need for specification of the positions along the sequence of the catalytic residues containing them. To generate the peptide binding cleft, we employed a version of RFD2 recently optimized for both enzyme and protein-protein interaction design, termed RFD2 for molecular interfaces (RFD2-MI)^10^, as the de novo design of proteases requires solving both challenges. We further anticipated that a cysteine-based catalytic triad mechanism could be advantageous for protease design, as this multistep mechanism has been extensively studied, catalytic triad based cleavage of activated esters has recently been demonstrated, and cysteine is more readily activated than serine. We set out to explore the use of RFD2-MI to design new cysteine proteases from scratch.

## Results

The first step in de novo enzyme design is the specification of a minimalist active site geometry. The cysteine protease mechanism involves a catalytic triad consisting of cysteine that serves as the nucleophile, an adjacent hydrogen-bonded histidine that activates the cysteine, stabilizes the leaving group, and promotes activation of water for deacylation, and an aspartate that hydrogen bonds to the histidine, tuning its p*K*_a_ and ensuring population of a catalytically competent tautomer. In addition, an oxyanion hole formed by backbone and/or sidechain hydrogen bond donors stabilizes the developing negative charge on the carbonyl oxygen along the reaction coordinate (Extended Data Fig. 1a). We chose to employ an oxyanion hole consisting of a backbone N-H group and a sidechain amide group. To position the substrate, we reasoned that β-strand pairing between the substrate, and the designed enzyme could enable precise placement of the amide bond to be cleaved relative to the catalytic machinery and compensate for the energetic cost of desolvating the polar amide groups along the substrate backbone. The cysteine protease Ulp1 (PDB 1EUV)^11^ has such an oxyanion hole, and binds substrates in a β-strand-like conformation stabilized by backbone hydrogen bonds; hence we used it as the basis for a minimal active site consisting of the catalytic triad (Ulp1 residues Cys580, His514, Asp531), the oxyanion hole (Gln574 and the backbones of 578–579), and four residues of the substrate backbone (Fig. 1a).

**Fig. 1.**
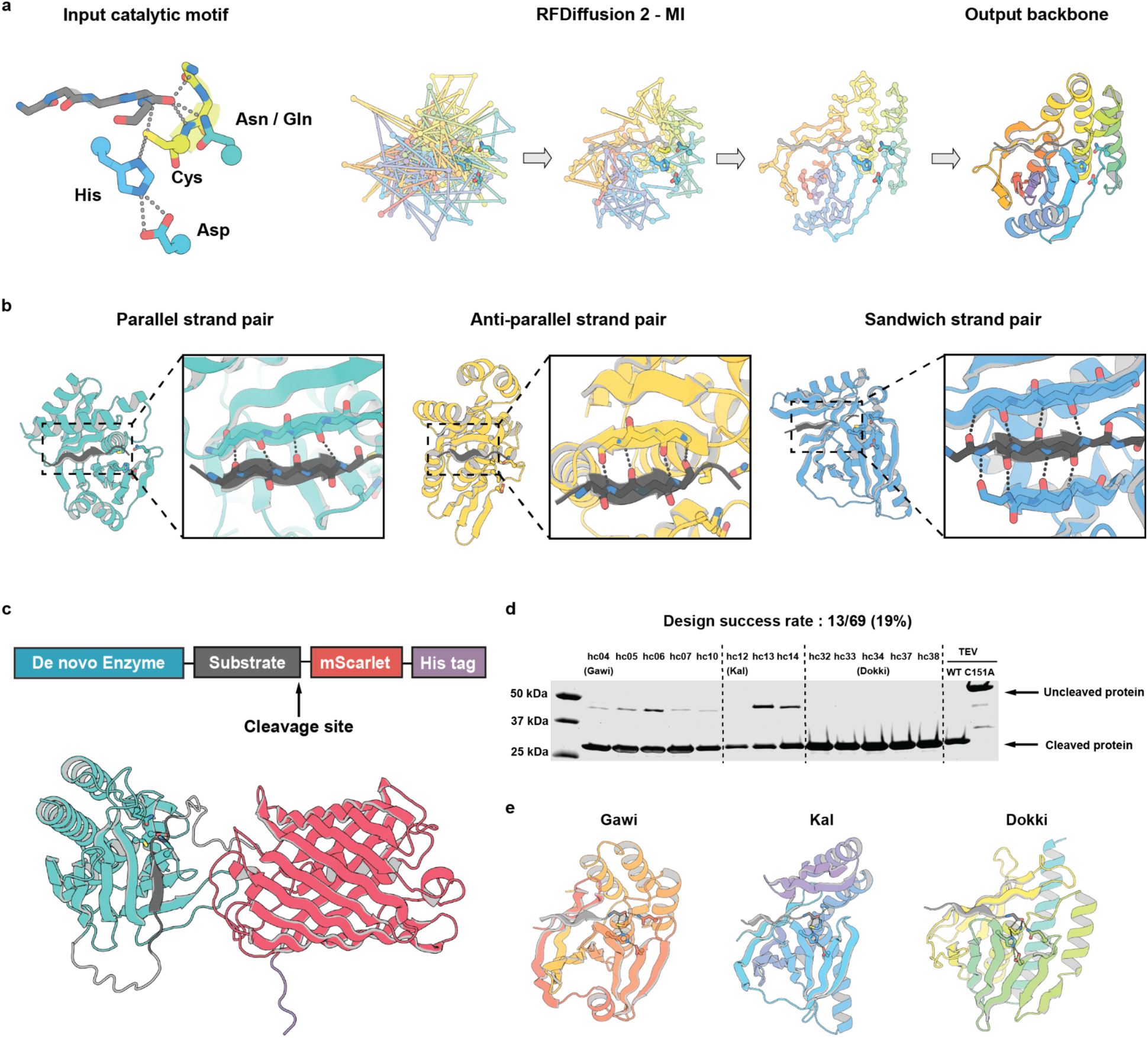
Computational design and experimental testing approach. **a**, Cysteine protease input catalytic triad motif from the Ulp1 crystal structure (PDB 1EUV) and backbone generation with RFDiffusion2-MI. **b**, Representative strand-paired binding-domain backbones from the computational design set. Structures were selected from the 3,232 backbones that passed DSSP computational filtering. **c**, Screening construct: the designed protease is co-expressed with a fusion substrate containing the target cleavage motif followed by mScarlet and a C-terminal His tag. Cleavage at the engineered scissile bond releases mScarlet-His. **d**, Post-IMAC, activity is verified by SDS–PAGE: active designs show a smaller mScarlet-His₆ peak/band, inactive designs show the larger full-length band. **e**, AF3 model of the Gawi, Kal, and Dokki.

To generate protein structures scaffolding this motif along with an appropriately placed peptide substrate, we used RFD2-MI (Fig. 1a). As we aimed to design a protease, but were agnostic about the exact sequence being cleaved, we chose to allow both the conformation and sequence of the substrate to be freely sampled in each design calculation, along with the designed enzyme. Starting from fully random placements of the enzyme and peptide substrate residues (Gaussian noise), RFD2-MI generated a wide range of outputs, many of which formed strand-pairing interactions with the substrate in parallel and antiparallel configurations (Fig. 1b). Sequences were designed for the resulting backbones with ProteinMPNN^12^, simultaneously optimizing the sequences of the enzyme and substrate to promote substrate binding and selectivity. Since protein substrates require relatively large binding sites which are susceptible to collapse in the absence of substrate, iterative design rounds were carried out in which the designs from the first two rounds were filtered for self-consistency based on AlphaFold 3^13^ apo-state predictions, retaining only those with RMSD < 1.2 Å over the designed active site. Designs for which AF3 models for both apo and complex states were very close (Cα RMSD < 0.8 Å, pLDDT > 90, and appropriate catalytic geometry; see Methods) to the design models (Extended Data Fig. 1b) were selected for experimental characterization.

To simplify screening, we fused the cognate substrate to the C-terminus of each design and appended a downstream mScarlet protein for detection^14^ and a hexahistidine tag for purification (Fig.1c). In this format, active designs undergo cleavage during *E. coli* expression, releasing free mScarlet, which is collected by nickel-affinity purification and then analyzed by SDS-PAGE and size-exclusion chromatography (SEC) (see Methods). Of the 69 designs tested, 13 exhibited cleavage activity (design success rate: 19%), corresponding to 3 of 12 unique backbones tested (Fig. 1d, Extended Data Fig. 2a,b). The most active designs from each of the three successful backbones were designated Gawi, Kal, and Dokki, respectively (Fig. 1d,e). Dokki and Kal exhibited complete substrate cleavage, whereas Gawi reached ∼95% cleavage (Fig. 1d). LC-MS analysis of SEC-purified protein samples confirmed that all 13 active designs cleaved precisely at the intended target site (Extended Data Fig. 3a, Supplementary table 2).

We tested the importance of the designed cysteine nucleophile, histidine base, and glutamine/asparagine oxyanion contact by mutating each residue individually to alanine and analyzing cleavage efficiency by SEC. Mutation of Cys or His to Ala abolished activity, confirming their essential roles in catalysis. By contrast, mutation of the Gln/Asn sidechain in the oxyanion hole did not uniformly inactivate the designs; in its absence, ordered water molecules may provide compensatory stabilization of the oxyanion intermediates (Extended Data Fig. 3b).

To determine if the designs could catalyze peptide bond cleavage in trans, we incubated the 13 designs that showed activity in the cis (fusion) assay with synthetic 8-residue substrate peptides labeled with an EDANS donor fluorophore and a Dabcyl quencher (Fig. 2a)^15^. The Gawi family exhibited very low activity on the FRET substrates, precluding kinetic analysis, and the Kal family could not be reliably characterized owing to poor substrate solubility. By contrast, the Dokki family showed measurable activity (Extended Data Fig. 4a,b). LC–MS analysis of the reaction products further confirmed precise cleavage at the intended target site of the Dokki cognate substrate (Extended Data Fig. 4c). The observed rates of the designs were modest (estimated *k*_cat_ values of ∼10^-6^ s^-1^ for Gawi and ∼10^-4^ s^-1^ for Dokki), so we pursued computational redesign of these two representative backbone groups using ProteinMPNN to enhance activity (see Methods; Extended Data Fig. 5a,b). In total, 21 redesigns were generated for Gawi and 89 for Dokki, of which 7 and 24, respectively, displayed increased activity, ranging from 1.2- to 31-fold faster than their parent designs (Extended Data Fig. 5c). For the best redesigned variants from each backbone, substitution of the catalytic Cys, His, or Asp residues with Ala significantly reduced activity, whereas mutation of the oxyanion hole residue reduced activity more modestly, yielding a 1.9- to 3.8-fold decrease (Fig. 2c), together confirming dependence on the designed active site. Steady-state kinetics revealed catalytic efficiencies (*k*_cat_/*K*_M_) of 0.24 ± 0.03 M^-1^ s^-1^ (*k*_cat_ = (0.8 ± 0.1) × 10^-5^ s^-1^, *K*_M_= 34 ± 4 μM) for Gawi-09 and 12 ± 2 M^-1^ s^-1^ (*k*_cat_ = (1.1 ± 0.1) × 10^-3^ s^-1^, *K*_M_ = 89 ± 10 μM) for Dokki-15 (Fig. 2d).

**Fig. 2.**
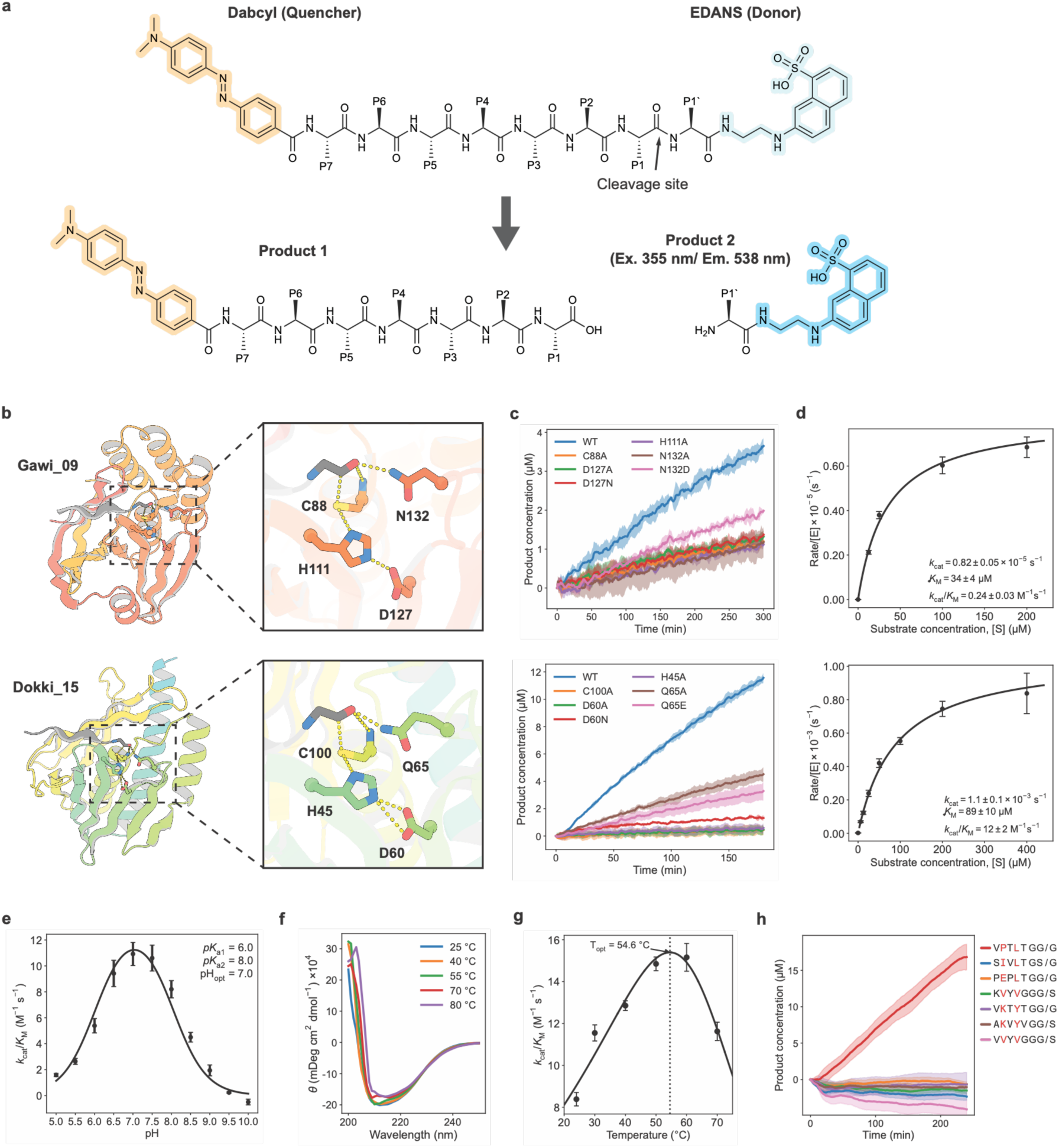
Designed proteases are active. **a,** Chemical structure of the Dabcyl–peptide–EDANS FRET substrate. **b,** Structural model of a representative designed protease showing the catalytic Cys–His–Asp triad and Asn/Gln oxyanion-hole residues. **c,** FRET cleavage assays comparing wild-type proteases with the corresponding active-site knockout variants. **d,**Steady-state kinetic measurements fitted to the Michaelis–Menten equation to estimate apparent *k*_cat_, *K*_M_ and *k*_cat_/*K*_M_. **e,** pH–rate profile of Dokki-15, showing a bell-shaped dependence with maximal activity at approximately pH 7.0. **f,** Thermal unfolding of Dokki-15 measured by circular dichroism, showing a melting temperature (*T*_m_) above 80 °C. **g,** Temperature–rate profile of Dokki-15, showing maximal activity at 54.6 °C and measurable activity at approximately 70 °C. **h,** FRET cleavage assays of Dokki-15 against diverse peptide substrates, demonstrating preferential cleavage of the intended substrate. For c–e, g and h, lines and shaded bands represent the mean and s.d., respectively, of three technical replicates (*n* = 3).

We pursued further biochemical characterization of the most efficient design, Dokki-15. The pH-rate profile revealed a bell-shaped curve with an apparent optimum at pH 7.0 (p*K*_a1_ = 6.0, p*K*_a2_ = 8.0) (Fig. 2e, Extended Data Fig. 6a,f). Like natural cysteine proteases, Dokki-15 shows a pH dependence consistent with ionization of the catalytic cysteine and histidine, which mediate nucleophilic attack and proton transfer, respectively. The apparent p*K*_a_ of the cysteine is shifted downward relative to the free thiol (typical p*K*_a_ ∼8-8.5, observed p*K*_a1_ = 6.0), consistent with stabilization of the thiolate by the neighboring histidine, but not as strongly shifted as in some native enzymes where cysteine p*K*_a_ values as low as 2.5-4.0 have been reported^16^. Similarly, the histidine p*K*_a_ is perturbed higher than the typical value in solution (typical p*K*_a_≈ 6, observed p*K*_a2_ = 8), favoring its charged form during the rate-limiting step. In many natural cysteine proteases, the resting state is described as a thiolate–imidazolium ion pair, supported by computational and experimental studies showing coupled Cys–His ionization and substantial p*K*_a_ perturbation^17^. While dyad protonation can be context-dependent, Dokki-15 reproduces the characteristic qualitative ionization pattern, albeit with more moderate p*K*_a_ shifts. Further optimization of electrostatic interactions could enhance the degree of cysteine activation and improve catalytic efficiency.

We assessed the stability of Dokki-15 using circular dichroism and found it to be highly thermostable, with a melting temperature (*T*_m_) above 80 °C (Fig. 2f). Dokki-15 retains enzymatic activity up to ∼70 °C, indicating that the designed catalytic triad remains functional even under conditions that typically disrupt protein folding and weaken enzyme-substrate interactions^18^ (Fig. 2g).

Substrate specificity was determined using a panel of eGFP–peptide–mScarlet fusion substrates. Dokki-15 exhibited high selectivity for its cognate substrate, VPTLTGG/G, with minimal or no detectable cleavage of seven non-cognate peptides sharing similar amphiphilic sequence features (Fig. 2h). Although several non-cognate substrates contained hydrophobic residues at positions corresponding to the P4-P6 region (highlighted in red), they were not efficiently cleaved, indicating that Dokki-15 has sequence specificity beyond general hydrophobic patterning. Thus, Dokki-15 specifically recognizes and cleaves its target substrate without exhibiting significant promiscuity toward related peptides.

We determined six X-ray crystal structures to assess the structural accuracy of the design method: Kal and Kal_C103A in complex with substrate, the apo structure of Gawi-09, the apo structure of Dokki, and both the apo and substrate-bound complex of Dokki-15. These span all three backbone scaffolds, and superposition with design models showed excellent agreement (Cα RMSDs < 1.2 Å), demonstrating the accuracy of the design method (Fig. 3, Supplementary Table 3). In the substrate-bound Kal complex, both the catalytic triad and substrate-binding cleft adopt the designed conformations (Fig. 3a). In the apo structure of Gawi-09, catalytic residues align closely with the model except for a displacement of His, likely caused by a Zn^2+^ ion from the crystallization buffer that bridges the catalytic His and Asp (Fig. 3b). For the apo structures of Dokki and Dokki-15, electron density consistent with a cysteine sulfonic acid indicates oxidation of the catalytic cysteine and disruption of the designed Cys–His interaction; in Dokki-15, this oxidized, substrate-free state is associated with displacement of the His-bearing β-hairpin and a 1.3-Å displacement of the His Cα atom (Fig. 3c–e). A similar oxidation-induced misalignment of His is observed in the native cysteine protease Ulp1^19^ (Extended Data Fig. 7e). Together, these structures confirm that the designed proteases adopt the intended folds and active-site geometries with near-atomic fidelity and function as designed.

**Fig. 3.**
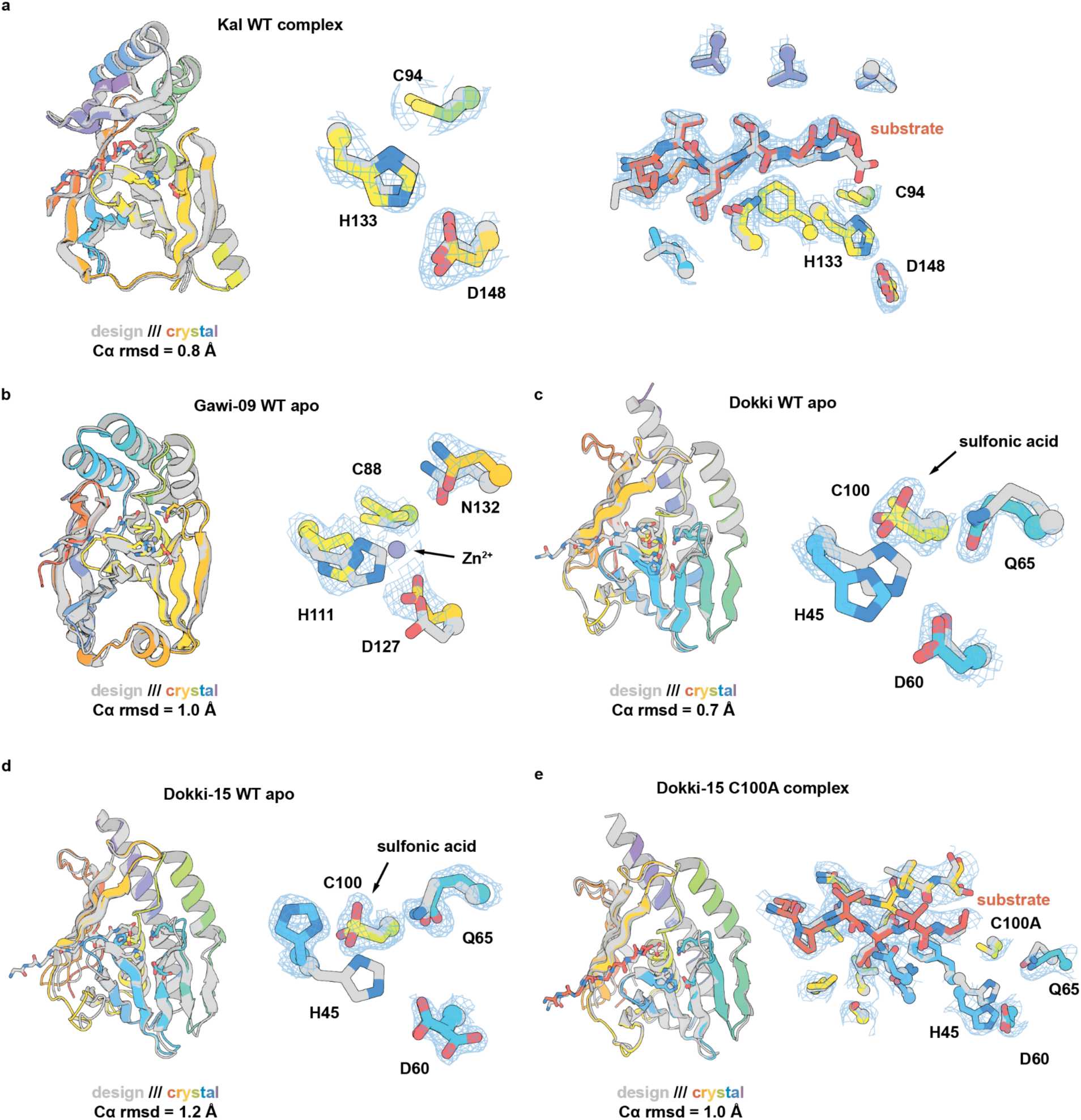
Structural characterization of designed cysteine proteases. **a**, Structural superposition of the design model (gray) and crystal structure (rainbow) for Kal WT complex, showing overlays of the catalytic and substrate-binding sites, resolution = 2.19Å. **b**, Superposition of design model (gray) and crystal structure (rainbow) for Gawi-09 WT apo, highlighting the catalytic site, resolution = 2.19Å. **c**, Superposition of design model (gray) and crystal structure (rainbow) for Dokki WT apo, highlighting the catalytic site, resolution = 1.77Å. **d**, Superposition of design model (gray) and crystal structure (rainbow) for Dokki-15 WT apo, highlighting the catalytic site, resolution = 1.96Å. **e**, Superposition of design model (gray) and crystal structure (rainbow) for Dokki-15 C100A complex, showing the substrate-binding site, resolution = 3.38Å.

To evaluate whether the designed proteases have novel structures and folds, we performed FoldSeek searches of the PDB and AlphaFold-derived datasets^20^. The lowest TM-score observed was 0.46 for Gawi-09, with all scaffolds scoring below 0.50—the threshold typically used to indicate fold similarity^21^ (Supplementary table 4). Structural alignments of these top hits revealed they have clearly distinct architectures, with backbone RMSDs exceeding several angstroms (Extended Data Fig. 7 a–d). These findings demonstrate that the designed proteases do not replicate existing natural folds but instead expand into previously unexplored regions of protein structure space, highlighting the innovative nature of our generative design approach, and sharply contrasting with previous efforts at redesigning native enzyme specificity.

To probe the origins of the remaining gap in activity between our designs and native cysteine proteases, we examined the effects of single amino acid substitutions at 20 positions surrounding the Dokki-15 active site that were predicted not to perturb the overall fold or catalytic geometry (Extended Data Fig. 8a,b; see Methods). We tested 185 single-site variants using the FRET-based assay (Extended Data Fig. 8c). Most substitutions introduced in the mutational screen markedly reduced enzyme activity (Fig. 4a), indicating that our original MPNN designed sequence was close to optimal for catalytic function. However, at several positions, Tyr41, Ser44, Pro67 and Thr99, beneficial substitutions were identified, none of which had been sampled in the original design calculations, as they were assigned low ProteinMPNN probabilities (Fig. 4a,b; Extended Data Fig. 8d). Combinations of these substitutions led to further activity increases, although variants containing Y41T, S44V, and T99F exhibited moderate substrate inhibition (Extended Data Fig. 8g–i). The most active variant, Dokki-v2 (Y41T:S44V:P67V:T99F), had a catalytic efficiency (*k*_cat_/*K*_M_) of 413 ± 50 M^-1^ s^-1^ (*k*_cat_= 0.064 ± 0.005 s^-1^; *K*_M_ = 154 ± 14 µM) (Fig. 4d; Extended Data Fig. 8j), and achieved more than 130 turnovers per active site. (Extended Data Fig. 8k). The improvement relative to Dokki-15 arises entirely from a 58-fold increase in the catalytic rate constant (*k*_cat_), which reports directly on the intrinsic chemical step at the active site. When this optimized *k*_cat_ value is compared with the reported rate constant for uncatalyzed peptide bond hydrolysis (*k*_uncat_ = 3.6 × 10^-11^ s^-1^ at 25 °C, pH 6.8^1^), Dokki-v2 is found to accelerate amide cleavage by an extraordinary 1.8 × 10^9^-fold.

**Fig. 4.**
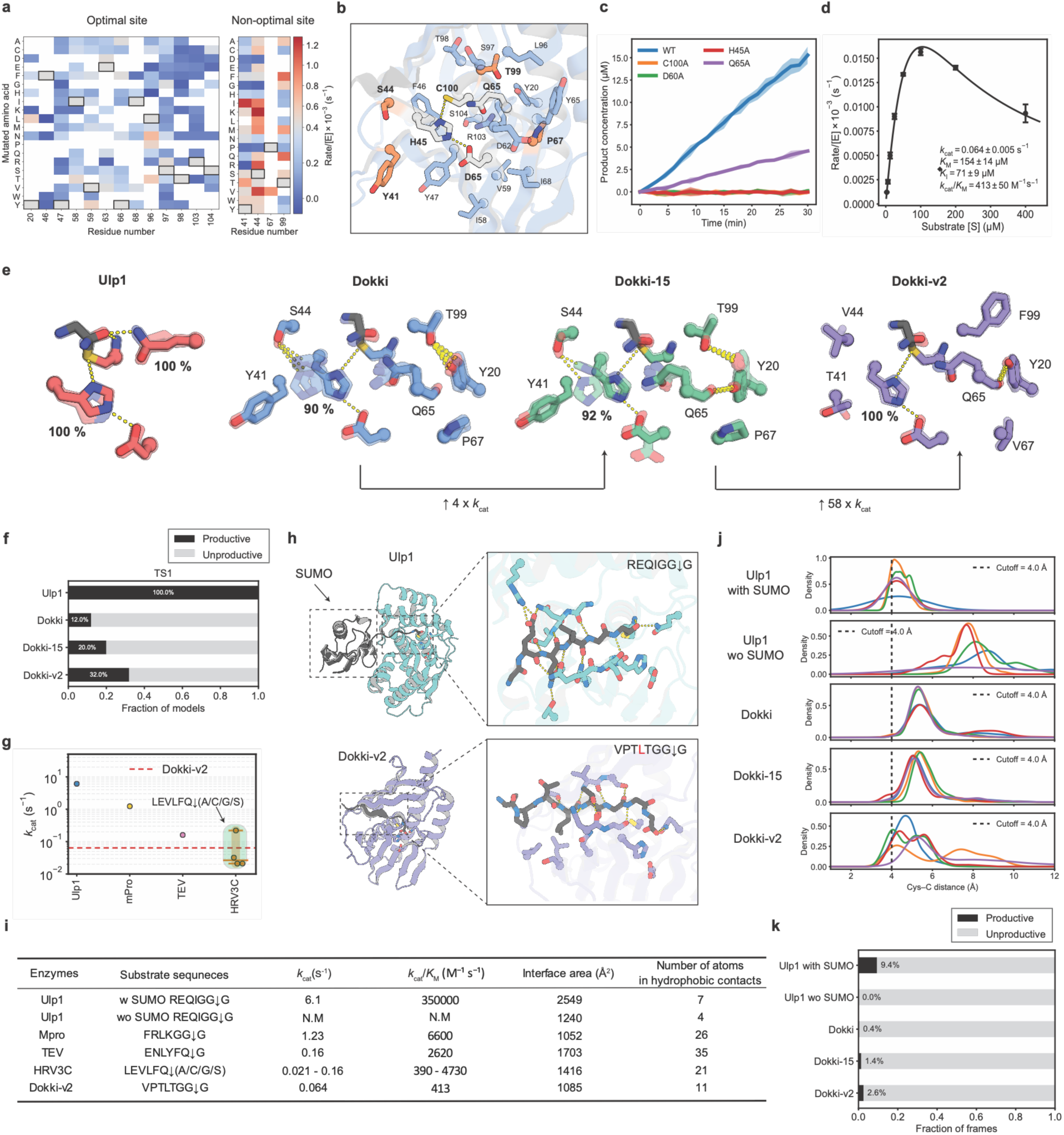
Contributions of active-site preorganization and substrate positioning to catalysis. **a,** Heat map of single-mutant activity measured with 3 µM enzyme and 75 µM substrate, reported as initial rate/[E] (s⁻¹). Red and blue indicate increased and reduced activity, respectively, relative to Dokki-15; white substitutions disrupted active-site organization *in silico* and were not tested. **b,** Activity-enhancing positions (Tyr41, Ser44, Pro67 and Thr99) mapped onto Dokki-15; catalytic residues are grey. **c,** FRET cleavage assays comparing wild-type designs and active-site knockout variants using 1 µM enzyme and 50 µM substrate. Lines and shading show the mean and s.d., respectively, of three technical replicates (*n* = 3). **d,** Steady-state kinetics of Dokki-v2 (Y41T:S44V:P67V:T99F), fitted to a Michaelis–Menten substrate-inhibition model to estimate apparent *k*_cat_, *K*_M_, *K*_I_ and *k*_cat_/*K*_M_. Data are mean ± s.d. of three technical replicates (*n* = 3). **e,** Representative TS1-state PLACER ensembles showing the catalytic triad, oxyanion-hole Gln and activity-enhancing sites. In Dokki and Dokki-15, His45 samples a non-productive interaction with Ser44. In Dokki-v2, Tyr20 instead contacts the Gln65 backbone carbonyl, stabilizing the oxyanion hole. **f,** Fraction of productive TS1-state PLACER conformations for Ulp1 and the Dokki series, defined by Cys–His <4.2 Å, His–Asp <3.2 Å and oxyanion-hole distance <3.4 Å. **g,** Reported turnover numbers (*k*_cat_) of representative sequence-specific native cysteine proteases and Dokki-v2. Ulp1 and Dokki-v2 were measured in-house using three technical replicates per enzyme (*n* = 3). TEV, Mpro and HRV3C values are literature-reported: TEV measurements were performed in duplicate per substrate concentration and Mpro assays in triplicate; HRV3C replication was not specified in the main article. HRV3C shading spans the minimum–maximum estimates across four substrates, LEVLFQ↓X (X = A, C, G or S). **h,** Substrate-binding pockets of Ulp1 and Dokki-v2; hydrophobic pocket residues are red. **i,** Catalytic and structural metrics, including *k*_cat_, *k*_cat_/*K*_M_, interface area and hydrophobic atom contacts calculated using PyRosetta. **j,** Distributions of catalytic Cys S–scissile carbonyl C distances from five independent 500-ns molecular-dynamics simulations per system (n = 5); traces denote individual replicas. **k,** Fraction of productive conformations sampled during the molecular-dynamics simulations, defined using the criteria in the main text.

To identify the structural basis of the activity gains, we analysed ensembles spanning the catalytic cycle of Ulp1 and the Dokki series using PLACER^22^, including the first transition state (TS1), the acyl–enzyme intermediate (AEI), the second transition state (TS2), and the apo states (Extended Data Fig. 9a–c). Productive conformations were defined using geometry thresholds derived from native cysteine proteases^23–29^ (Cys–His < 4.2 Å, His–Asp < 3.2 Å and oxyanion-Gln < 3.4 Å; Extended Data Fig. 9d). TS1 most clearly separated Ulp1 from the Dokki designs and best tracked activity, consistent with initial nucleophilic attack being rate-limiting. In Dokki and Dokki-15, His45 samples a non-productive rotamer stabilized by Ser44; the Y41T and S44V substitutions in Dokki-v2 suppress this state, improving dyad preorganization. P67V and T99F enhance positioning of Gln65 for oxyanion stabilization via a Tyr20-mediated hydrogen bond to its backbone carbonyl (Fig. 4e). The oxyanion-Gln45 distance is still longer than in Ulp1 (Extended Data Fig. 9b), resulting in fewer productive TS1 conformations (36% versus 100%; Fig. 4f), consistent with the modest effect of the Gln knockout and a limited contribution of this interaction to catalysis (Fig. 4c).

To benchmark activity, we compared Dokki-v2 with sequence-specific native cysteine proteases (Ulp1, Mpro, TEV, and HRV3C)^30–32^ (Fig. 4g and Supplementary Table 5). The *k*_cat_ value for Dokki-v2 falls at the lower end of the native range, approximately one to two orders of magnitude below the most active native enzymes. Structural comparison shows that Mpro, TEV, and HRV3C proteases form well-defined substrate-binding interfaces, which utilize key hydrophobic contacts to position the scissile bond (Fig. 4h; Extended Data Fig. 9h). By contrast, Ulp1 engages the folded SUMO domain via a large binding surface to position its C-terminus containing the scissile bond in the active site. Our designs utilize the catalytic triad and oxyanion hole geometry of Ulp1, albeit on entirely different protein scaffolds, but do not contain an additional auxiliary folded domain positioning the substrate in the active site. For a direct comparison, we measured Ulp1 activity on peptide substrates lacking the SUMO domain (Supplementary Fig.2). We found that while Ulp1 cleaves SUMO-containing substrates (*k*_cat_ ∼ 6.1 s^-1^), it shows no detectable activity with eGFP–peptide–mScarlet substrates lacking the SUMO domain (Supplementary Fig.2). Interface analysis indicates that, although Ulp1 has fewer local hydrophobic contacts, its larger binding surface compensates via extended interactions (Fig. 4i). In contrast, Dokki-v2 forms a relatively smaller interface with limited hydrophobic contacts compared to native protease, suggesting weaker substrate engagement.

Molecular dynamics (MD) simulations^33^ of the Michaelis complex (5 x 500 ns replicas) support the interpretation that precision of positioning of the substrate amide bond to be cleaved relative to the catalytic cysteine accounts for at least part of the remaining observed activity differences to the native enzymes (Fig. 4j and Extended Data Fig. 9f–g). For Ulp1 with the peptide-only substrate, productive ES conformations (Cys–C < 4.0 Å, Cys–His < 4.2 Å and His–Asp < 3.2 Å) were rarely sampled due to dissociation from the binding pocket, consistent with the lack of activity (Fig. 4j–k). For Dokki-v2, productive conformations were sampled more frequently, consistent with the observed activity, but at a lower rate than Ulp1 on the SUMO containing substrate, primarily owing to more effective positioning of the scissile carbonyl relative to the catalytic cysteine. These results suggest that the short and relatively polar peptide substrate of Dokki-series provides weaker substrate engagement than the native cysteine proteases, limiting its ability to enforce productive scissile-bond presentation, and that higher activity could be achievable on targets with larger hydrophobic residues as in Mpro, TEV, and HRV3C, or with additional positioning domains as in the Ulp1-SUMO case.

Lastly, we investigated whether the computationally designed proteases are active in living mammalian cells using a nuclear-localized fluorescence reporter system. The designed proteases fused to sfGFP and an mScarlet reporter were directed to chromatin through fusion to histone H2B. The reporter contains the protease substrate sequence between the sfGFP and mScarlet (Fig. 5a,b), and hence protease-mediated cleavage can be detected by release and redistribution of the mScarlet signal from the nucleus. Cells expressing the Dokki series, including the original Dokki designs straight out of the computer, showed clear relocalization of mScarlet away from the nucleus, whereas cells expressing the inactive Cys-to-Ala mutants retained nuclear colocalization of sfGFP and mScarlet (Fig. 5c–h; Extended Data Fig. 10a–g; Supplementary Fig.3). These results demonstrate that the de novo designed proteases function in mammalian cells, creating opportunities for a wide range of synthetic biology and cellular engineering applications^34–37^.

**Fig. 5.**
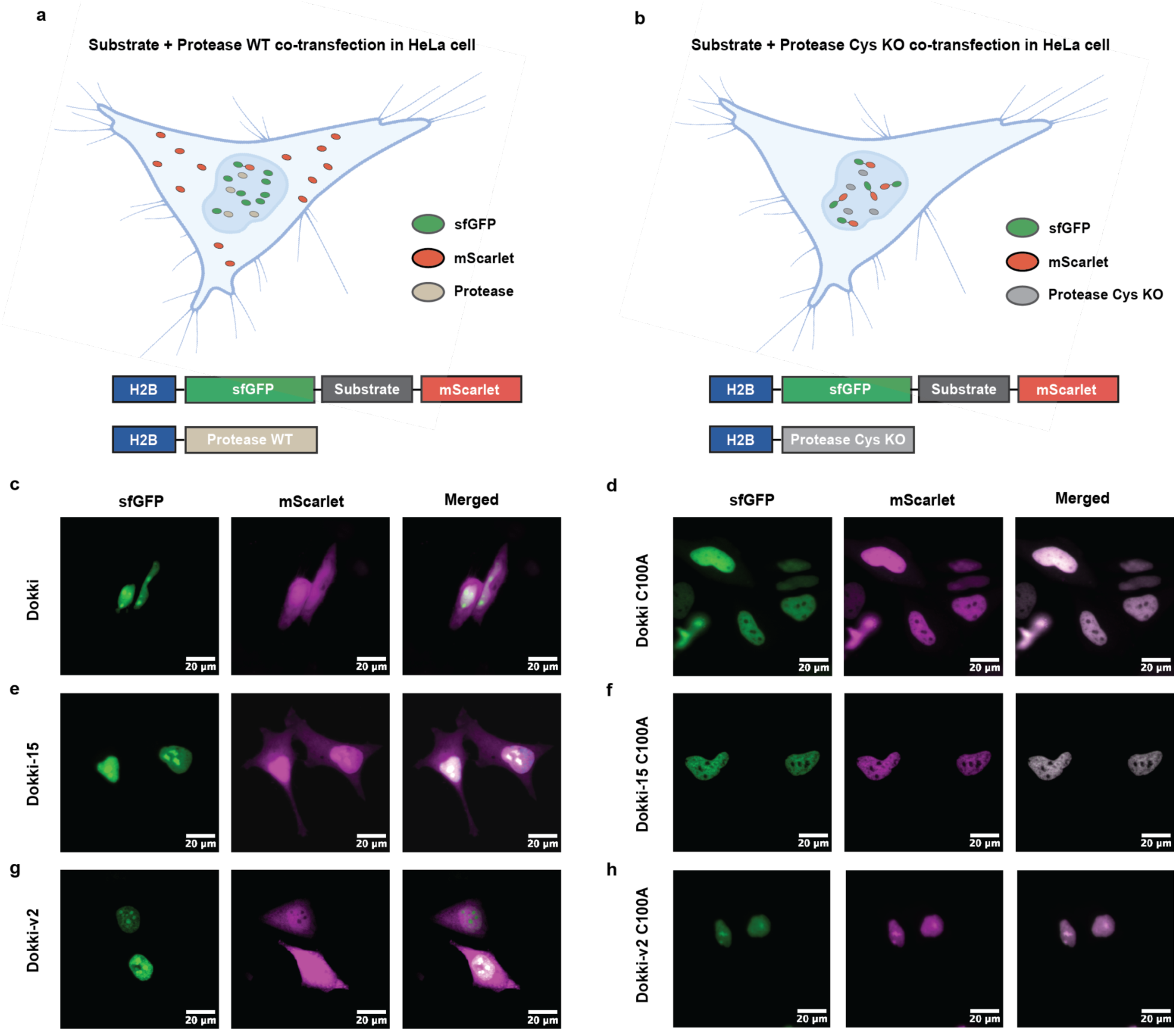
Designed proteases function in mammalian cells. **a,b,** Schematic of the reporter assay. HeLa cells were co-transfected with plasmids encoding H2B–sfGFP–substrate–mScarlet and an H2B-fused protease or its catalytic Cys-to-Ala knockout variant. H2B directs the fusion proteins to the nucleus. Protease-mediated substrate cleavage releases mScarlet from the nuclear H2B–sfGFP reporter, resulting in redistribution of the mScarlet signal. Schematics in **a,b** were created with BioRender.com. **c–h,** Representative fluorescence microscopy images of HeLa cells showing sfGFP, mScarlet and merged channels at the indicated time after transfection. Cells were transfected with substrate and protease plasmids at a 100-ng:10-ng ratio and imaged by InCell microscopy. Images are representative of 3 independent transfection experiments with similar results. **c,** Dokki; **d,** Dokki C100A; **e,** Dokki-15; **f,** Dokki-15 C100A; **g,** Dokki-v2; **h,** Dokki-v2 C100A.

## Discussion

While previous computational efforts have largely focused on chemically simple or artificially activated model reactions, our results here demonstrate that generative design can now tackle one of the most energetically demanding transformations in biology—the hydrolysis of amide bonds within protein substrates. The most active designs prior to optimization achieve rate enhancements of up to 3 × 10^7^ over the uncatalyzed reaction, underscoring the catalytic potential of de novo–designed enzymes. The designed enzymes depend on the Cys–His–Asp catalytic triad, cleave precisely at the intended peptide bond, function in living mammalian cells, and exhibit atomic-level structural fidelity to design models (Cα RMSDs <1.2 Å) while adopting folds distinct from natural proteases (TM-scores < 0.5), together demonstrating both the precision and creative breadth of the design process.

Despite these advances, catalytic efficiency remains at the lower end of native cysteine protease performance (Fig. 4g, Supplementary table 5). Our analyses suggest that there is room for improvement both in the precise geometry of the catalytic machinery, and in positioning the peptide substrate relative to this machinery so the peptide bond to be cleaved is properly oriented relative to the catalytic cysteine. In papain-like native cysteine proteases, the Cys–His–Asp triad and oxyanion hole have been evolutionarily tuned to achieve highly optimized geometric alignment^38^ (Extended Data Fig. 9d). Our probing of the sequence dependence of catalysis suggests that despite the successful recapitulation of active site geometry evident in the crystal structures of our designs, the pre-organization of the active site is still suboptimal, and subtle errors in positioning may account for much of the remaining gap in activity. For example, PLACER ensembles of the most active variants suggest that substitutions at positions 41 and 44 in Dokki-15 improve catalytic-dyad preorganization by suppressing non-productive histidine rotamers, and substitutions at positions 67 and 99 stabilize the oxyanion-hole glutamine. Incorporating these preorganization-enhancing substitutions yielded our top-performing design, which has a *k*_cat_ = 0.064 s^-1^ and *k*_cat_/*K*_M_ = 413 M^-1^ s^-1^, only ∼2.5-fold and ∼6-fold lower than the widely used native cysteine protease TEV (*k*_cat_= 0.16 s^-1^; *k*_cat_/*K*_M_= 2620 M^-1^ s^-1^), respectively^30^. The remaining activity gap relative to Ulp1, which we used as a template for the catalytic geometry, likely reflects both imperfect oxyanion-hole placement and suboptimal substrate alignment with the catalytic machinery. Consistent with this interpretation, MD simulations and structural analyses indicate that the peptide substrate is less precisely oriented relative to the catalytic cysteine in the designed enzymes compared to native cysteine proteases on their native substrates, and we find that the designs have much greater activity than Ulp1 has on a comparably sized peptide substrate (lacking the folded SUMO domain) which simulations suggest is not well positioned in the Ulp1 active site.

There are several routes forward to further improve the design methodology and obtain still more active catalysts. First, as noted above, methodological improvements should prioritize increasing active-site preorganization, particularly with respect to catalytic triad alignment and oxyanion stabilization. Ensemble-focused optimization, for example with PLACER, can help identify catalytic residues to target to restrict mobility and disfavor phenomena like the observed non-productive histidine rotamers. Going beyond fixed backbone sequence design could help identify such interactions as several of the activity increasing substitutions had very low ProteinMPNN sequence probabilities; our in silico single substitution screen which allows small backbone adjustments and does identify these activity increasing substitutions could for example be incorporated into the original computational design pipeline. Lastly, while the present work modeled only a subset of the substrate (P1-P7), extending the design to include a larger binding interface and additional hydrophobic interactions could enhance substrate complementarity, increase *k*_cat_, and enable specificity toward diverse target sequences. Such improvements should enable the design of increasingly efficient and selective cysteine proteases which could have transformative potential in biotechnology and therapeutic applications.

The relatively high success rate of our computational design methodology, together with the finding that the designed proteases cleave their substrates in mammalian cells, opens the door to exciting new synthetic biology and biomedical applications. For example, because computationally designed proteases can be generated for a wide variety of target sequences, synthetic signaling cascades^34^ can expand considerably beyond what is possible with native proteases, and designed proteases targeting endogenous disease-relevant substrates could have direct therapeutic utility.

## Methods

### Computational design of cysteine proteases

#### RFDiffusion2 for Molecular Interface (RFD2-MI)

We performed backbone generation using RFD2-MI in three rounds: an initial round of diffusion from complete noise followed by two rounds of iterative partial diffusion (T = 40 and T = 10). Catalytic residues (Cys580, His514, Asp531), oxyanion hole residues (Gln574, backbones 578–579), and substrate residues (chain B, 95–98) were extracted from the Ulp1 crystal structure (PDB: 1EUV) and used as the input motif. By permuting motif order, we generated enzyme backbones of 180–200 residues paired with a 7-residue substrate segment. From 50,000 initial backbones, structures were filtered using DSSP analysis to exclude those with excessive loops^39^ (>8 consecutive residues), low strand content (<0.1), or high coil content (>0.3), yielding 3,232 candidates. After one diffusion–ProteinMPNN–AlphaFold3 (apo prediction) cycle, 104 parent backbones that passed AF3 filters were subjected to a round of partial diffusion (T = 40), each producing 150 variants. DSSP filtering yielded 6,402 backbones, of which 208 advanced through the ProteinMPNN–AF3 cycle. These were used as input for a final round of partial diffusion (T = 10), generating 30 variants per backbone.

#### Sequence design

Filtered backbones were subjected to 14, 39, and 24 cycles of ProteinMPNN^12^ and Rosetta FastRelax^40^ in rounds one, two, and three, respectively. To preserve catalytic geometry, interatomic distances were constrained with 0.1 Å tolerances: NE2(His)–OD2(Asp), 2.8 Å; NE2(His)–OD1(Asp), 3.2 Å; ND1(His)–SG(Cys), 3.6 Å; ND1(His)–N(P1′), 4.4 Å; O(P1)–ND2(Asn), 3.0 Å; C(P1)–SG(Cys), 3.1 Å. Catalytic triad residues (Cys, His, Asp) and the oxyanion hole residue (Asn/Gln) were fixed during design, and Met, Cys, and His were disallowed during design.

#### Filtering

In the first two rounds, only apo structures were predicted with AF3^13^. For the first round, designs were retained if they satisfied pLDDT > 80, pTM > 0.80, and Cα RMSD to the MPNN model < 1.8 Å. For the second round, the criteria were tightened to pLDDT > 85, pTM > 0.85, and Cα RMSD < 1.2 Å. In the final round, both apo and complex predictions were performed, and filtering incorporated structural confidence, catalytic geometry, and complex stability. Designs were retained if they met the following criteria: complex pLDDT > 90, ipTM > 0.90, pae_min < 1.2, and key_plddt > 85; catalytic distances (NE2(His)–OD1(Asp) < 3.2 Å, ND1(His)–SG(Cys) < 3.8 Å, SG(Cys)–C < 3.5 Å, with ≥2 oxyanion contacts); and apo criteria (pLDDT > 90, Cα RMSD to complex prediction model < 0.8 Å).

#### Computational re-design using ProteinMPNN guided by AF3

To improve activity, we focused optimization on two backbones with soluble and testable FRET substrates: Gawi backbone, which showed minimal activity but potential for improvement, and Dokki backbone, which contained the most active design. Redesign was performed using a pLDDT-guided strategy: (i) residues with pLDDT < 85 were resampled with ProteinMPNN and retained only if their pLDDT improved to > 90, and (ii) residues within 4 Å of the substrate were redesigned to strengthen binding interactions. All variants were subsequently filtered for high-confidence, atomically accurate AF3 predictions of the enzyme-substrate complex (Extended Data Fig. 5a,b).

### Design and screening of enzyme-substrate fusion constructs

#### DNA cloning, expression and protein purification for screening

The gene sequences for the individual designs were codon-optimized for expression in *Escherichia coli*. Linear DNA fragments (eBlocks; Integrated DNA Technologies) with overhangs compatible with Golden Gate cloning were inserted into the pCOOL6 vector (GenScript), a modified expression plasmid containing a kanamycin resistance cassette and a cloning cassette (MSG–BsaI–ccdB–BsaI–(GS)\₄–mScarlet–GS–6×His). Golden Gate assembly reactions (5 µL) were set up in 96-well PCR plates and incubated at 37 °C for 1 h. Subsequently, 2 µL of each reaction mixture was used to transform 10 µL of chemically competent *E. coli* BL21(DE3) cells. Cells were recovered in 100 µL S.O.C. medium for 2 h and grown overnight in 1 mL LB containing 50 µg mL^-1^ kanamycin. For protein expression, 10 µL of overnight culture was inoculated into 1 mL LB and shaken at 1,300 rpm, 37 °C for 2 h, followed by induction with 10 µL 100 mM IPTG and overnight (16hr) culture at 37 °C. Cultures were harvested by centrifugation (4,000g, 5 min), and pellets were resuspended in 400 µL lysis buffer (20 mM HEPES pH 7.0, 300 mM NaCl, 25 mM imidazole, 1× BugBuster). After 30 min shaking at 1,300 rpm, 25 °C, lysates were clarified (4,300g, 15 min) and supernatants applied to Ni^2+^-charged MagBeads (L00295, GenScript). Bound proteins were washed with buffer (20 mM HEPES pH 7.0, 300 mM NaCl, 25 mM imidazole) and eluted with buffer (20 mM HEPES pH 7.0, 300 mM NaCl, 500 mM imidazole). Eluates were filtered through a 96-well 0.22-µm filter plate (203940-100, Agilent) prior to analysis.

#### Screening for proteolytic activity by SDS–PAGE, SEC, and Liquid chromatography–mass spectrometry (LC–MS)

For SDS–PAGE, 15 µL of each eluate was mixed with 5 µL of 4× Laemmli buffer, heated to 95 °C for 5 min, and resolved on Bio-Rad 26 wells AnykD Criterion precast gels. Gels were stained with Coomassie blue, destained, and imaged using a LI-COR Odyssey M imager. Uncleaved constructs (enzyme–substrate–mScarlet fusion) migrated at ∼48 kDa, whereas cleaved mScarlet appeared at ∼27 kDa. In parallel, eluates were analyzed by size-exclusion chromatography (SEC) on a Superdex 75 Increase 10/300 GL column (Cytiva) equilibrated in 20 mM HEPES pH 7.0, 150 mM NaCl. Elution was monitored at 520 nm to track mScarlet with 0.65 mL/min. Uncleaved proteins eluted at ∼2.0 mL, whereas cleaved mScarlet eluted at ∼2.5 mL. SEC fractions containing cleaved mScarlet were analyzed by reverse-phase LC-MS using an Agilent G6230B TOF instrument fitted with an AdvanceBio RP-Desalting column. Spectra were deconvoluted with Bioconfirm software employing a total entropy algorithm. Each backbone group encoded a distinct substrate sequence, leading to backbone-specific cleavage products. In all cases, observed molecular weights corresponded to theoretical values, confirming precise cleavage at the intended recognition site.

### Kinetic characterization of protease designs with soluble FRET substrates

#### DNA cloning, expression and protein purification for enzyme characterization

Genes encoding selected hits were subcloned into the LM1371 expression plasmid (MS-6×His-BsaI-ccdB-BsaI) to generate enzyme constructs lacking the covalently attached substrate and mScarlet used during screening. In this format, each designed protease was expressed as a free enzyme with an N-terminal hexahistidine tag in *E. coli*BL21(DE3) cells, enabling purification by Ni–NTA chromatography. For protein production, 500 µL of overnight culture was used to inoculate 50 mL LB medium supplemented with 50 µg mL^-1^ kanamycin, followed by incubation at 37 °C with shaking (225 rpm) for 2 h. Expression was induced with 50 µL of 1 M IPTG, and cultures were incubated overnight at 37 °C. Cells were harvested by centrifugation (4,200g, 5 min), resuspended in 35 mL lysis buffer (20 mM HEPES pH 7.0, 300 mM NaCl, 25 mM imidazole, 1 mM TCEP), and sonicated on ice water (10 min total, 80% amplitude, 5 s on/5 s off). Resulting lysates were clarified by centrifugation (14,000g, 30 min), and the supernatant was loaded onto 1 mL Ni-NTA resin equilibrated with wash buffer. Resin was washed three times with 20 mL wash buffer and proteins were eluted with 5 mL elution buffer (20 mM HEPES pH 7.0, 300 mM NaCl, 500 mM imidazole, 1 mM TCEP). Elution fractions were further purified by size-exclusion chromatography (Superdex 75 Increase 10/300 GL, Cytiva) in running buffer (20 mM HEPES pH 7.0, 150 mM NaCl). Purified proteins were used immediately for downstream assays or flash frozen in liquid nitrogen and stored at −80 °C. Protein molecular weights were verified by liquid chromatography–mass spectrometry (LC–MS).

#### FRET assay

Fluorogenic substrates were synthesized by GenScript with a DABCYL quencher at the N terminus, a protease-specific recognition sequence, and an EDANS donor at the C terminus, separated by GS or GSGS linkers^41^. Each substrate was mass-verified by GenScript using MS prior to use (>95% purity by HPLC). Substrate stock solutions were prepared by dissolving solid in DMSO to a concentration of 10 mM. For kinetic characterization, purified enzyme was incubated with FRET substrate in 20 µL reaction volumes in low-volume 384-well plates (Corning). Reactions were prepared in assay buffer (20 mM HEPES pH 7.0, 150 mM NaCl, 14 mM β-mercaptoethanol, 5% DMSO). Proteolysis was monitored in real time at 37 °C using a Neo2 plate reader (excitation 355 nm, emission 538 nm, gain 100). Fluorescence was calibrated by measuring RFU values for a dilution series of free EDANS (50, 25, 12.5, 6.25, 3.125, 1.5625, and 0 µM). The data were fit to a linear regression, yielding the conversion factor: RFU=311.56×[EDANS], where [EDANS] is the free fluorophore concentration in µM (Extended Data Fig. 6a). Fluorescence increase corresponding to EDANS release was analyzed by nonlinear regression with a custom Python script to extract catalytic parameters (*k*_cat_ and *K*_M_). Initial rates (*v*_0_, µM·s^-1^) were obtained from the initial slope of product formation after converting RFU to µM using a calibration factor.

When reported as apparent first-order rate constants, we normalized by the total enzyme concentration to give *k*_obs_ = *v*_0_/[*E*]_0_ (s⁻¹). These values were fit to the Michaelis–Menten equation: *k*_obs_=*k*_cat_[S]/(*K*_M_+[S]) where [S] is the substrate concentration, *k*_cat_ is the catalytic turnover number, and *K*_M_ is the Michaelis constant. For variants exhibiting substrate inhibition, data were instead fit to a substrate inhibition model: *v*=*V*_max_[S]/(*K*_M_+[S](1+[S]/*K*_I_)) where *K*_I_ represents the apparent inhibition constant describing formation of an unproductive enzyme–substrate complex.

#### pH-rate profile

To assess the pH-dependence of activity, a set of buffers were prepared: 20 mM sodium acetate (pH 5.0), 20 mM MES (pH 5.5–6.5), 20 mM HEPES (pH 7.0–7.5), 20 mM Tris (pH 8.0–8.5), and 20 mM CHES (pH 9.0–10.0), each supplemented with 150 mM NaCl and 14 mM BME. FRET assays were performed as described above in triplicate at each pH. FRET assays were performed in triplicate at an enzyme concentration of 10 µM and a substrate concentration of 10 µM. RFU values were converted to EDANS concentration using pH-specific calibration factors (RFU = calibration factor × [EDANS]µM; pH 5.0: 316.49, pH 5.5: 344.55, pH 6.0: 320.67, pH 6.5: 315.09, pH 7.0: 311.56, pH 7.5: 306.10, pH 8.0: 316.84, pH 8.5: 300.42, pH 9.0: 271.20, pH 9.5: 252.32, pH 10.0: 224.29; Extended Data Fig. 6a).

Initial rates were extracted from background-corrected fluorescence time courses: replicate traces were smoothed with a centered rolling mean (window = 6 points), the no-enzyme control was subtracted, negative values were clipped at zero, and the first 60 min were fit by linear regression to obtain initial velocities (µM s^-1^). Slopes were converted to apparent specificity constants by dividing by enzyme concentration and substrate concentration using the low-substrate approximation *v*_0_≈(*k*_cat_*/K*_M_)[E][S] (Extended Data Fig. 6f). Reported values are thus *k*_cat_*/K*_M_ with units of M^-1^ s^-1^, calculated as mean ± s.d. across replicates for each pH. The pH–rate profile was fit by nonlinear least squares to a two-ionization model, *v*_0_ = V_max_/(1+10^p*K*1−pH^+10^pH−p*K*2^), with V_max_, p*K*_1_, and p*K*_2_ free. When replicate standard deviations were available, they were used as weights (otherwise the fit was unweighted); parameters are reported with 95% confidence intervals from the covariance matrix. The apparent optimum pH was computed as pH_opt_=(p*K*_1_+p*K*_2_)/2.

#### Temperature-rate profile

To assess activity across temperatures, FRET assays were performed in triplicate at each condition using 8 µM enzyme and 10 µM substrate. Fluorescence values (RFU) were converted to EDANS concentrations using the calibration factor RFU = 311.56 × [EDANS]. Initial rates were determined from background-corrected fluorescence time courses: replicate traces were smoothed with a centered rolling mean (window = 6 points), the no-enzyme control was subtracted, negative values were clipped to zero, and the first 30 min were fit by linear regression to obtain initial velocities (µM s^-1^). Rates were converted to catalytic efficiencies (*k*_cat_/*K*_M_) by dividing by enzyme and substrate concentration and using the low-substrate approximation *v*₀ ≈ (*k*_cat_/*K*_M_)[E][S]. Reported values are expressed as *k*_cat_/*K*_M_ (M^-1^ s^-1^) (Extended Data Fig. 6g).

#### Substrate specificity assays

To assess substrate specificity, peptide substrates were expressed as fusion proteins of the form MSG–eGFP–peptide–mScarlet–6×His. Constructs were generated by subcloning peptide sequences into the pCOOL6 expression plasmid and expressed in *E. coli* BL21(DE3) cells. Substrates were purified using the same expression and purification procedures described for enzyme production.

The following peptide sequences were tested: VPTLTGG/G (Dokki-15 cognate substrate), SIVLTGS/G, PEPLTGG/G, KVYVGGG/S, VKTYTGG/G, AKVYVGG/S, VVYVGGG/S

Proteolytic activity toward each substrate was measured by a FRET-based assay. Reactions were initiated by mixing enzyme and substrate solutions at a 1:1 ratio to yield final concentrations of 5 µM enzyme and 37.5 µM substrate (prepared from 10 µM enzyme and 75 µM substrate stocks). Fluorescence was monitored over a 4 h time course, and product formation was quantified by normalizing fluorescence values to the signal obtained from fully cleaved substrate using a calibration factor of 4262 RFU per 37.5 µM product.

#### Total turnover number determination

Total turnover number (TTN) was measured over 24 h using FRET assays performed in triplicate with 10 µM enzymes for Dokki-15 and 1 µM enzyme for Dokki-v2 and substrate concentrations of 400, 200, 100, or 50 µM. Fluorescence values (RFU) were converted to EDANS concentrations using the calibration factor RFU = 311.56 × [EDANS]. TTN was calculated by dividing the total product concentration by the enzyme concentration (Extended Data Fig. 6h, 8k).

#### Circular dichroism (CD) spectroscopy

Secondary structure content was assessed by circular dichroism (CD) spectroscopy using a Jasco J-1500 instrument equipped with a Peltier-based temperature control system (EXOS). Measurements were carried out in quartz cuvettes with a 0.1 cm optical path length over a wavelength range of 200–250 nm. Thermal unfolding was monitored by recording changes in ellipticity at 222 nm while heating from 25 to 80 °C at a rate of 1 °C min^-1^. In addition, full spectra were collected at 25, 40, 55, 70, and 80 °C. CD data were expressed as molar ellipticity (θ), with mean residue ellipticity values calculated using a protein concentration of 10 µM (0.2 mg/mL).

### Site-saturation mutagenesis

#### In silico and wet-lab site-saturation mutagenesis

Residues located within 4.5 Å of the Dokki-15 catalytic residues (Cys100, His45, Asp60, and Gln65) were selected for site-saturation mutagenesis. Each position was individually mutated to the remaining 19 amino acids, generating a library of single-mutation variants. For each variant, enzyme–substrate complexes were modeled using AF3. Variants were retained if they met the following thresholds: complex pLDDT > 92 (wild type, 95.3), minimum PAE < 0.9 (wild type, 0.795), inter-chain predicted TM-score (ipTM) > 0.93 (wild type, 0.95), and key-residue pLDDT > 93 (wild type, 96.3). In total, 185 single-mutation variants satisfied these criteria and were selected for experimental characterization (Extended Data Fig. 8a,b).

All variants were expressed in *E. coli* BL21(DE3) cells and purified by IMAC followed by SEC. Protein concentrations were determined by UV absorbance and normalized to 6 µM prior to activity measurements. Enzymatic activity was assessed by a FRET assay, in which enzyme and peptide substrate solutions were mixed at a 1:1 ratio to yield final concentrations of 3 µM enzyme and 75 µM substrates. Initial reaction rates were calculated and normalized to the wild-type enzyme. Activities of all single-mutation variants were visualized as a heat map (Extended Data Fig. 8c). Mutations exhibiting enhanced activity were subsequently combined to generate multi-mutation variants, which were expressed, purified, and evaluated under identical assay conditions.

### Computational analysis

#### PLACER modeling and catalytic geometry analysis

To analyze the active-site geometry in the tetrahedral intermediate states, ensembles of PLACER models were generated using 50 independent random seeds, yielding 50 models per design. AF3-predicted enzyme–substrate complex structures served as input for PLACER. A covalent bond was introduced between the substrate carbonyl carbon atom (C) and the catalytic cysteine sulfur (SG) to mimic the tetrahedral intermediate.

For each model within each ensemble, distances describing the oxyanion hole and catalytic geometry were measured. These distributions were compared across Ulp1 PLACER models, non-active designs, Dokki-15, and Dokki-v2. Native reference geometries were derived from a curated set of crystal structures (PDB IDs: 1TL9, 1ATK, 8A4V, 1NQC, 1ME3, 3BPF, 1BP4, 1CVZ, 1PPN, and 1EUV).

#### Molecular dynamics simulations

MD simulations of the Michaelis complex were performed with the GPU-accelerated version of AMBER24 (pmemd.cuda). Protein atoms were modeled with the ff14SB force field, and systems were prepared in tleap. Each complex was solvated in a truncated octahedral box of TIP3P water with an 8 Å buffer, and Na^+^ and Cl^−^ ions were added to neutralize the system. Energy minimization was carried out in two stages of 5,000 steps each (2,500 steepest descent followed by 2,500 conjugate gradient): first with harmonic restraints of 500 kcal mol^−1^ Å^−2^ applied to all non-hydrogen protein atoms, excluding solvent and ions, and then with restraints of 50 kcal mol^−1^ Å^−2^ applied to backbone heavy atoms only. Systems were heated from 0 K to 300 K over 300 ps under NVT conditions with a 1 fs time step and backbone restraints of 5 kcal mol^−1^ Å^−2^, using Langevin dynamics for temperature control (collision frequency, 5.0 ps^−1^). This was followed by 50 ns of equilibration at 300 K and 1 atm under NPT conditions with a 2 fs time step and a Monte Carlo barostat (relaxation time, 10.0 ps). Bonds involving hydrogen atoms were constrained using SHAKE, and long-range electrostatics were treated with particle mesh Ewald using an 8 Å direct-space cutoff. Production simulations were performed as five independent 500-ns replicas under NPT conditions at 300 K and 1 atm, using a Monte Carlo barostat relaxation time of 5.0 ps. Coordinates were saved every 200 ps and analyzed with cpptraj after removal of solvent and ions and RMSD fitting to the minimized structure.

#### Interface surface-area and hydrophobic-contact analysis

Protein–substrate interface properties were analyzed using PyRosetta. For each input PDB structure, chains A and B were parsed as the two interacting partners. Interface metrics were calculated using the Rosetta InterfaceAnalyzerMover with the interface defined as A_B and separated-state packing enabled. The extracted metrics included interface buried solvent-accessible surface area (interface_delta_sasa), that was used as the Rosetta-reported buried interface surface area between the two chains.

Hydrophobic contacts across the chain A–chain B interface were quantified independently from the Rosetta interface metrics. Hydrophobic residues were defined as Ala, Val, Leu, Ile, Met, Phe, Trp, Tyr and Pro. For each pair of hydrophobic residues across the two chains, all heavy atoms were examined, and hydrophobic atoms were approximated as carbon. Pairwise distances were calculated between all hydrophobic atoms from the two residues. A residue–residue hydrophobic contact was counted once if at least one hydrophobic atom pair was within 4.0 Å. For each structure, the analysis returned the total number of hydrophobic atom–atom contacts.

### Kinetic characterization of Ulp1 protease with eGFP-SUMO-peptide-mScarlet substrates

#### DNA cloning, protein expression and purification

The gene encoding Ulp1 was subcloned into the LM1371 expression plasmid, MS-6×His-BsaI-ccdB-BsaI. Substrate constructs containing either SUMO–REQIGGG or REQIGGG alone were subcloned into the pCOOL6 expression plasmid and expressed as fusion proteins of the form MSG–eGFP–peptide–mScarlet–6×His. Each construct was transformed into *E. coli* BL21(DE3) cells for protein expression. For protein production, 5 mL of overnight culture was used to inoculate 500 mL of LB medium supplemented with 50 µg mL^-1^ kanamycin. Cultures were grown at 37 °C with shaking at 225 rpm for 2 h, after which protein expression was induced by addition of 500 µL of 1 M IPTG. Cultures were then incubated overnight at 20 °C. Cells were harvested and proteins were purified by immobilized metal-affinity chromatography followed by size-exclusion chromatography. Purified proteins were concentrated, and protein concentrations were determined by absorbance using a NanoDrop spectrophotometer

#### Gel-based kinetic characterization

Ulp1 activity against eGFP–peptide–mScarlet substrates were measured using a gel-based cleavage assay. For substrates lacking SUMO, reactions contained 1 µM Ulp1 and substrate concentrations generated by twofold serial dilution from 90 to 2.8 µM. Reactions were incubated for 1 h at room temperature. For SUMO-containing substrates, reactions contained 30 nM Ulp1 and substrate concentrations generated by twofold serial dilution from 20 to 0.6 µM. Reactions were incubated for 1 min at room temperature.

Following incubation, reaction products were analyzed by SDS–PAGE. Gels were stained with Coomassie blue and imaged using a Li-COR imaging system with Coomassie detection. Band intensities were quantified in ImageJ, and the fraction cleaved was calculated from the relative intensities of substrate and product bands. Initial rates were determined from the cleaved fraction, substrate concentration and reaction time, and kinetic parameters were obtained by fitting the data to the Michaelis–Menten equation. All measurements were performed in three replicates.

### Mammalian cell assays

#### DNA preparation for expression in HeLa cells

Gene sequences encoding individual designs were codon-optimized for human cell expression. Linear DNA fragments (eBlocks; Integrated DNA Technologies) containing overhangs compatible with Golden Gate cloning were assembled into the ZJ101 vector, a modified expression plasmid carrying an ampicillin-resistance cassette. Golden Gate assembly reactions were performed in 5-µL volumes in 96-well PCR plates at 37 °C for 1 h. For transformation, 2 µL of each assembly reaction was added to 10 µl chemically competent *E. coli* DH5α(DE3) cells. Cells were recovered in 100 µL S.O.C. medium for 2 h, plated on LB agar containing 50 µg mL^-1^ ampicillin, and incubated overnight at 37 °C. Plasmids confirmed by DNA sequencing were used for subsequent experiments.

#### Cell culture and transfection

HeLa cells were obtained from the American Type Culture Collect (ATCC). Authentication was provided by the vendor, ATCC. The cell lines were not tested for mycoplasma contamination. The cells we used were consistent with the size and shape of HeLa, and the doubling time and attachment durability were also consistent. 15,000 HeLa cells were seeded in 96 well glass bottom plate plates with #1.5 cover glass (Cellvis, P96-1.5H-N) in 100 uL DMEM (Gibco; Thermo Fisher Scientific) containing 10% Fetal Clone II Serum (Cytiva), 25 mM glucose, 1 mM sodium pyruvate, and 4 mM GlutaMAX. The following day, the cells were transfected with 13 µL of transfection mixture per well. The transfection mixture contained 10ng of the protease or protease knockout, 100ng of the substrate, 0.2 µL Lipofectamine 2000 (Thermo Fisher Scientific), and Opti-Mem Medium (Gibco) up to 13 µL. The mixture was incubated at room temperature for 30 minutes prior to adding to cells. The media was replaced with fresh FluoroBrite DMEM (Gibco) supplemented with 10% Fetal Clone II Serum 6 hours after transfection. The plates were prepared for imaging 24-48 hours after transfection as described in the subsequent section.

#### Hela cell imaging and analysis

Prior to imaging, the culture medium was exchanged for FluoroBrite DMEM supplemented with NucBlue Live Cell DNA dye (Thermofisher, Cat# R37605), 1.5 drops per mL media, and cells were incubated at 37 °C for 15 min to permit labeling. Fluorescence images were automatically sampled using an IN Cell Analyzer 2500HS (Molecular Devices) and a 20× 0.75 NA CFI Plan Apo objective (Nikon). Cells were illuminated with a seven-color Solid State Illuminator (SSI) for fluorescence excitation. Fluorescence signals were acquired sequentially using the following filters: green (excitation 473 nm, emission 511/23 nm), red (excitation 575 nm, emission 623/24 nm), and blue (excitation 390 nm, emission 435/48 nm). Imaging was controlled using the IN Cell Analyzer 2500 HS software version 7.4 and light collected on sCMOS camera without binning. Nine fields of view were acquired for each well.

Images were background corrected using the rolling ball algorithm in Fiji (sigma=75). A custom Jupyter notebook was used to identify nuclei using the NucBlue signal by first applying a gaussian blur (sigma=1) followed by a triangle threshold from the scikit-image.filters package. Small objects were removed (max size=1500) and cells on the image edge were removed from analysis. Labeled images were created from the remaining nuclei, these objects were filtered to retain only cells that expressed GFP. Final object intensity was measured in both the red and green channels and the ratio of object mean green fluorescence/object mean red fluorescence was plotted for each condition. On average 200 cells were analyzed per condition. Ratios of green fluorescence/ red fluorescence were normalized based on a negative control sample.

### Structural characterization

#### Crystallography

Crystallization experiments were conducted using the sitting drop vapor diffusion method in 96-well plate format at 20 °C. Crystallization plates with 200 nL drops were set up using a Mosquito LCP from SPT Labtech, then imaged using UVEX microscopes and UVEX PS-256 from JAN Scientific. The Kal WT complex was crystallized in 2.8 M sodium acetate trihydrate pH 7.0. The Kal C103A complex was crystallized in 60% (v/v) tascimate pH 7.0. Dokki WT apo was crystallized in 1.6 M sodium citrate pH 6.5 and cryoprotected with 25% glycerol during harvest. Gawi-09 WT apo was crystallized in 0.05 M zinc acetate dihydrate, 0.05 M MES pH 6.1, and 11% (w/v) PEG 8,000, and cryoprotected with 25% PEG 8,000 during harvest. Dokki-15 WT apo was crystallized in 0.1 M BIS-TRIS pH 6.5, and 20% (w/v) polyethylene glycol monomethyl ether 5,000. Dokki-15 C100A complex was crystallized in 0.1 M citric acid pH 3.5, and 25% (w/v) PEG 3,350, and cryoprotected in 20% MPD during harvest. All diffraction quality crystals were frozen in liquid nitrogen for synchrotron data collection.

Diffraction data was collected either at the Advanced Photon Source beamline NE-CAT 24ID-E or National Synchrotron Light Source II, beamline 17-ID-1/17-ID-2. X-ray intensities and data reduction were evaluated and integrated using XDS^42^ and merged/scaled using Pointless/Aimless in the CCP4 program suite^43^. Structure determination and refinement starting phases were obtained by molecular replacement using Phaser^44^ using the designs as the search models. Following molecular replacement, the models were improved using Phenix^45^/Refmac^43^, and using simulated annealing. Structures were refined in Phenix ^45^/Refmac^43^. Model building was performed using COOT^46^. The final model was evaluated using MolProbity^47^. Data collection and refinement statistics are recorded in supplementary Table 3. Data deposition, atomic coordinates, and structure factors reported in this paper have been deposited in the Protein Data Bank (PDB), http://www.rcsb.org/ with accession code 9YNL (Dokki-15 C100A complex), 9YNM (Dokki-15 WT apo), 9YOX (Gawi-09 WT apo), 9YOY (Kal C103A complex), 9YOZ (Kal WT complex) and 9YP0 (Kal WT apo)

#### Structural similarity search

Structural similarity searches were carried out using Foldseek web server (https://search.foldseek.com/search) in TM-align mode against the AlphaFold/UniProt50 v6, AlphaFold/Swiss-Prot v6, AlphaFold/Proteome v6, PDB100 (20240101), BFVD 2023_02, CATH50 4.3.0, MGnify-ESM30 v1, BFMD 20240623, and GMGCL 2204 databases. For each design, the top structural hit and corresponding TM-score were recorded: Kal WT complex, 0.48 (AFDB50, AF-A0A923ZJ62-F1-model_v6); Kal C103A complex, 0.49 (AFDB50, AF-A0A923ZJ62-F1-model_v6); Dokki WT apo, 0.50 (all databases, A0A346A683); Gawi-09 WT apo, 0.46 (AFDB50, AF-A0A0X8G3I4-F1-model_v6); Dokki-15 WT apo, 0.50 (MGnify-ESM30, MGYP001077622663); Dokki-15 C100A complex, 0.48 (CATH50, 2.170.210.20 Q9N4L1 1–108).

#### Statistics and reproducibility

No statistical method was used to predetermine sample size. FRET measurements were performed using three technical replicates per condition. Cell-imaging experiments were performed in 3 independent transfection experiments, with nine fields acquired per well and 200 cells on average analyzed per condition. No data was excluded. No randomization or blinding was used.

## Data availability

Atomic coordinates and structure factors generated in this study are publicly available through the RCSB Protein Data Bank (https://www.rcsb.org/) under accession codes 9YNL, 9YNM, 9YOX, 9YOY, 9YOZ and 9YP0. Source data underlying Figs. 1–5 and Extended Data Figs. 1–10 are provided with this paper. Publicly available structures used as computational inputs or structural references were obtained from the RCSB Protein Data Bank under accession codes 1EUV, 1TL9, 1ATK, 8A4V, 1NQC, 1ME3, 3BPF, 1BP4, 1CVZ and 1PPN. Protein sequences are provided in Supplementary Table 1, and complete plasmid DNA sequences are provided in Supplementary Information 1 and 2.

## Code availability

Code used for computational design and analysis is publicly available at Github (https://github.com/hojaec/RFD2_MI_cysteine_protease_pipeline). Code used for biochemical kinetic data and mammalian-cell images analysis is publicly available at Github (https://github.com/hojaec/Cysteine-protease-data-analysis).

## Acknowledgments

Crystallographic data collection was performed on APS NE-CAT 24ID-E beamline and the beam time award(s) (https://doi.org/10.46936/APS-189869/60014006) from the Advanced Photon Source, a U.S. Department of Energy (DOE) Office of Science user facility operated for the DOE Office of Science by Argonne National Laboratory under Contract No. DE-AC02-06CH11357. This research used resources (FMX/AMX) of the National Synchrotron Light Source II, a U.S. Department of Energy (DOE) Office of Science User Facility operated for the DOE Office of Science by Brookhaven National Laboratory under Contract No. DE-SC0012704. The Center for BioMolecular Structure (CBMS) is primarily supported by the National Institutes of Health, National Institute of General Medical Sciences (NIGMS) through a Center Core P30 Grant (P30GM133893), and by the DOE Office of Biological and Environmental Research (KP1607011). Crystallographic data collected at NSLS II either under NECAT BAG proposal #317877 or WestCoastBAG2026. Schematic illustrations in Fig. 5 were created with BioRender.com.

## Author contributions

H.C. and S.J.P. developed the computational design pipeline, with contributions from P.V. B.C. and M.S.B. developed the RFD2-MI model. H.C. designed and characterized the proteases. S.S. and J.D. contributed to the mammalian-cell experiments. G.G., K.D., L.L.S. and G.R.L. contributed to the computational analyses. A.K.B., A.K., H.N., E.J., B.S. and S.J.P. determined and analyzed the crystal structures. D.H., S.J.P. and D.B. supervised the research. H.C., D.H., S.J.P. and D.B. wrote the manuscript. M.S.B., A.C., D.K., T.R.T., J.M.G. and A.F.S. contributed to the manuscript. All authors reviewed and commented on the manuscript.

## References

1. Radzicka, A. & Wolfenden, R. Rates of uncatalyzed peptide bond hydrolysis in neutral solution and the transition state affinities of proteases. J. Am. Chem. Soc. 118, 6105–6109 (1996).

2. Dyer, R. P. & Weiss, G. A. Making the cut with protease engineering. Cell Chem. Biol. 29, 177–190 (2022).

3. Otto, H.-H. & Schirmeister, T. Cysteine proteases and their inhibitors. Chem. Rev. 97, 133–172 (1997).

4. Carter, P. & Wells, J. A. Dissecting the catalytic triad of a serine protease. Nature 332, 564–568 (1988).

5. Ménard, R. et al. Modification of the electrostatic environment is tolerated in the oxyanion hole of the cysteine protease papain. Biochemistry 34, 464–471 (1995).

6. Verma, S., Dixit, R. & Pandey, K. C. Cysteine proteases: Modes of activation and future prospects as pharmacological targets. Front. Pharmacol. 7, 107 (2016).

7. Lauko, A. et al. Computational design of serine hydrolases. Science 388, eadu2454 (2025).

8. Kim, D., et al. Computational Design of Metallohydrolases. bioRxiv (2024) doi:10.1101/2024.11.13.623507.

9. Ahern, W. et al. Atom-level enzyme active site scaffolding using RFdiffusion2. Nat. Methods 23, 96–105 (2026).

10. Bauer, M. S., et al. *De novo* design of phospho-tyrosine peptide binders. bioRxiv (2025) doi:10.1101/2025.09.29.678898.

11. Mossessova, E. & Lima, C. D. Ulp1-SUMO crystal structure and genetic analysis reveal conserved interactions and a regulatory element essential for cell growth in yeast. Mol. Cell 5, 865–876 (2000).

12. Dauparas, J. et al. Robust deep learning-based protein sequence design using ProteinMPNN. Science 378, 49–56 (2022).

13. Abramson, J. et al. Accurate structure prediction of biomolecular interactions with AlphaFold 3. Nature 630, 493–500 (2024).

14. Bindels, D. S. et al. mScarlet: a bright monomeric red fluorescent protein for cellular imaging. Nat. Methods 14, 53–56 (2017).

15. Matayoshi, E. D., Wang, G. T., Krafft, G. A. & Erickson, J. Novel fluorogenic substrates for assaying retroviral proteases by resonance energy transfer. Science 247, 954–958 (1990).

16. Hofer, F., Kraml, J., Kahler, U., Kamenik, A. S. & Liedl, K. R. Catalytic site pKa values of aspartic, cysteine, and Serine proteases: Constant pH MD simulations. J. Chem. Inf. Model. 60, 3030–3042 (2020).

17. Zhai, X. & Meek, T. D. Catalytic mechanism of cruzain from Trypanosoma cruzi as determined from solvent kinetic isotope effects of steady-state and pre-steady-state kinetics. Biochemistry 57, 3176–3190 (2018).

18. Daniel, R. M. & Danson, M. J. A new understanding of how temperature affects the catalytic activity of enzymes. Trends Biochem. Sci. 35, 584–591 (2010).

19. Xu, Z. et al. Molecular basis of the redox regulation of SUMO proteases: a protective mechanism of intermolecular disulfide linkage against irreversible sulfhydryl oxidation. FASEB J. 22, 127–137 (2008).

20. van Kempen, M. et al. Fast and accurate protein structure search with Foldseek. Nat. Biotechnol. 42, 243–246 (2024).

21. Zhang, Y. & Skolnick, J. Scoring function for automated assessment of protein structure template quality. Proteins 57, 702–710 (2004).

22. Anishchenko, I. et al. Modeling protein-small molecule conformational ensembles with PLACER. Proc. Natl. Acad. Sci. U. S. A. 122, e2427161122 (2025).

23. LaLonde, J. M. et al. Use of papain as a model for the structure-based design of cathepsin K inhibitors: crystal structures of two papain-inhibitor complexes demonstrate binding to S’-subsites. J. Med. Chem. 41, 4567–4576 (1998).

24. Tsuge, H. et al. Inhibition mechanism of cathepsin L-specific inhibitors based on the crystal structure of papain-CLIK148 complex. Biochem. Biophys. Res. Commun. 266, 411–416 (1999).

25. Moldoveanu, T., Campbell, R. L., Cuerrier, D. & Davies, P. L. Crystal structures of calpain-E64 and -leupeptin inhibitor complexes reveal mobile loops gating the active site. J. Mol. Biol. 343, 1313–1326 (2004).

26. Falke, S. et al. Structural elucidation and antiviral activity of covalent cathepsin L inhibitors. J. Med. Chem. 67, 7048–7067 (2024).

27. Yamamoto, A. et al. Structural basis for development of cathepsin B-specific noncovalent-type inhibitor: crystal structure of cathepsin B-E64c complex. Biochim. Biophys. Acta 1597, 244–251 (2002).

28. Pauly, T. A. et al. Specificity determinants of human cathepsin s revealed by crystal structures of complexes. Biochemistry 42, 3203–3213 (2003).

29. Huang, L., Brinen, L. S. & Ellman, J. A. Crystal structures of reversible ketone-Based inhibitors of the cysteine protease cruzain. Bioorg. Med. Chem. 11, 21–29 (2003).

30. Kapust, R. B. et al. Tobacco etch virus protease: mechanism of autolysis and rational design of stable mutants with wild-type catalytic proficiency. Protein Eng. 14, 993–1000 (2001).

31. Kenneson, J. R. et al. Exploring possible drug-resistant variants of SARS-CoV-2 main protease (Mpro) with noncovalent preclinical candidate, Mpro61. ACS Bio Med Chem Au 5, 215–226 (2025).

32. Mei, M. et al. A combinatorial strategy for HRV 3C protease engineering to achieve the N-terminal free cleavage. Int. J. Biol. Macromol. 265, 131066 (2024).

33. Pearlman, D. A. et al. AMBER, a package of computer programs for applying molecular mechanics, normal mode analysis, molecular dynamics and free energy calculations to simulate the structural and energetic properties of molecules. Comput. Phys. Commun. 91, 1–41 (1995).

34. Gao, X. J., Chong, L. S., Kim, M. S. & Elowitz, M. B. Programmable protein circuits in living cells. Science 361, 1252–1258 (2018).

35. Roybal, K. T. et al. Engineering T cells with customized therapeutic response programs using synthetic Notch receptors. Cell 167, 419–432.e16 (2016).

36. Barnea, G. et al. The genetic design of signaling cascades to record receptor activation. Proc. Natl. Acad. Sci. U. S. A. 105, 64–69 (2008).

37. Coukos, R. et al. An engineered transcriptional reporter of protein localization identifies regulators of mitochondrial and ER membrane protein trafficking in high-throughput CRISPRi screens. Elife 10, e69142 (2021).

38. Novinec, M. & Lenarčič, B. Papain-like peptidases: structure, function, and evolution. Biomol. Concept*s***4**, 287–308 (2013).

39. Hekkelman, M. L., Salmoral, D. Á., Perrakis, A. & Joosten, R. P. DSSP 4: FAIR annotation of protein secondary structure. Protein Sci. 34, e70208 (2025).

40. Bennett, N. R. et al. Improving de novo protein binder design with deep learning. Nat. Commun. 14, 2625 (2023).

41. Kuo, C.-J., Chi, Y.-H., Hsu, J. T.-A. & Liang, P.-H. Characterization of SARS main protease and inhibitor assay using a fluorogenic substrate. Biochem. Biophys. Res. Commun. 318, 862–867 (2004).

42. Kabsch, W. Integration, scaling, space-group assignment and post-refinement. Acta Crystallogr. D Biol. Crystallogr. 66, 133–144 (2010).

43. Winn, M. D. et al. Overview of the CCP4 suite and current developments. Acta Crystallogr. D Biol. Crystallogr. 67, 235–242 (2011).

44. McCoy, A. J. et al. Phaser crystallographic software. J. Appl. Crystallogr. 40, 658–674 (2007).

45. Adams, P. D. et al. PHENIX: a comprehensive Python-based system for macromolecular structure solution. in International Tables for Crystallography 539–547 (International Union of Crystallography, Chester, England, 2012).

46. Emsley, P. & Cowtan, K. Coot: model-building tools for molecular graphics. Acta Crystallogr. D Biol. Crystallogr. 60, 2126–2132 (2004).

47. Williams, C. J. et al. MolProbity: More and better reference data for improved all-atom structure validation. Protein Sci. 27, 293–315 (2018).

