## Extended data figure 1-10, Supplementary table 1-5, Supplementary Fig 1-3, Supplementary information 1-2 for "De novo design of cysteine proteases"

#### Extended Data Figures

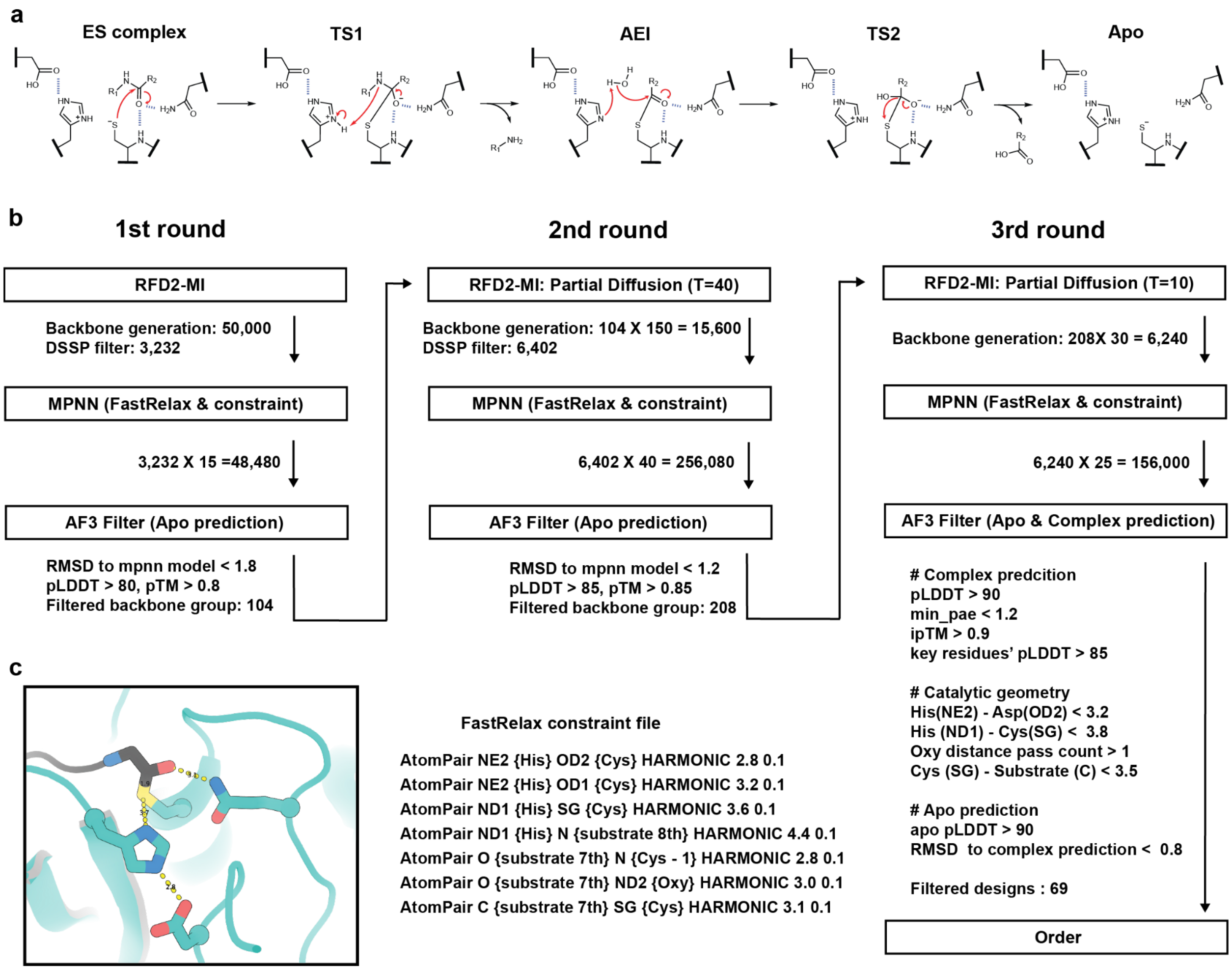

#### Extended Data Fig. 1 | Cysteine protease mechanism and design pipeline.

**a**, Mechanism of cysteine protease **b**, Overview of the computational pipeline: iterative cycle of (RFD2-MI → ProteinMPNN → AF3), with the number of designs retained at each step. **c**, Example of constraint file used in Rosetta FastRelax to preserve catalytic geometry during sequence optimization.

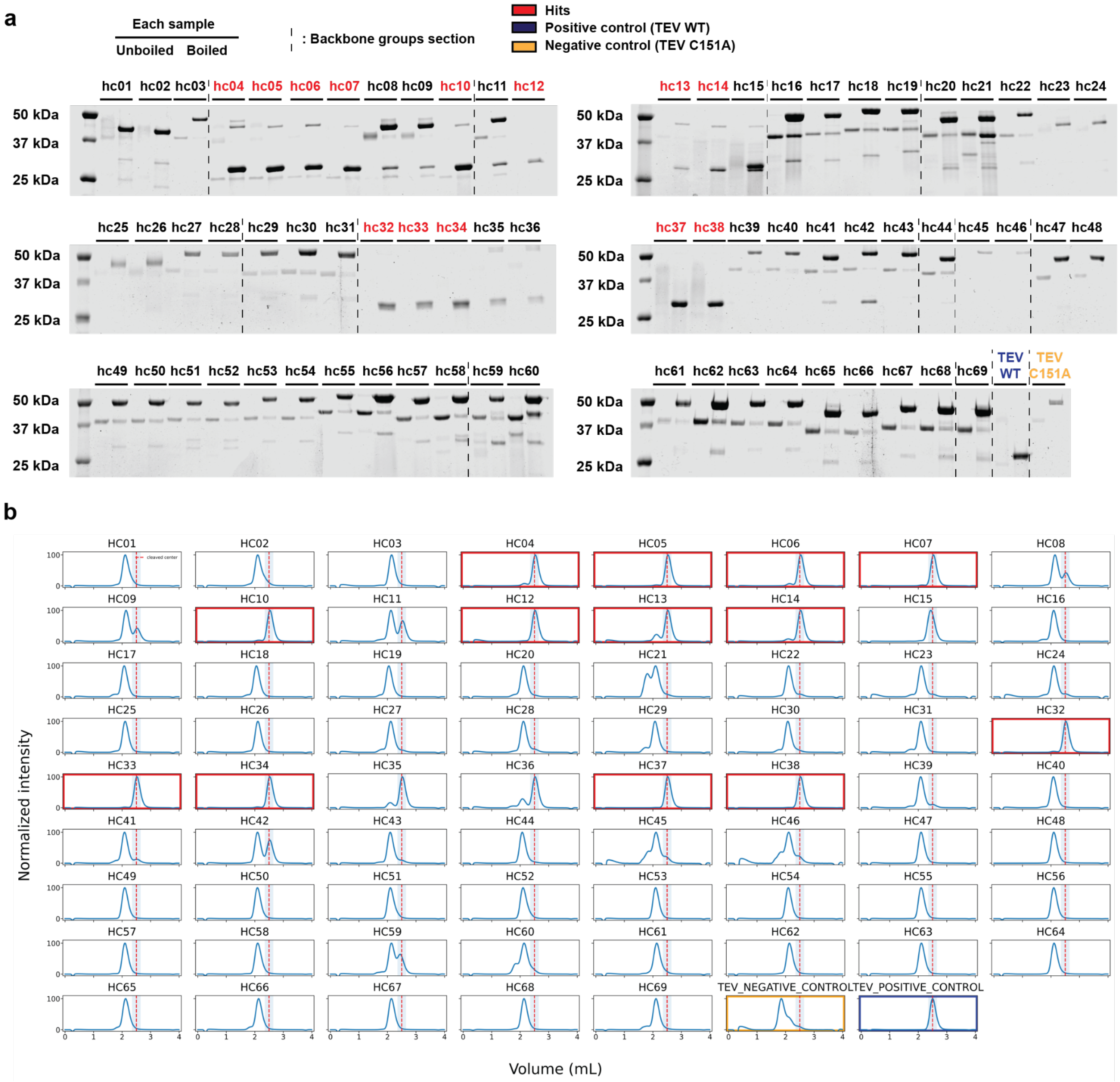

**Extended Data Fig. 2 | SDS-PAGE gel and HPLC–SEC screening of designed proteases.**

**a**, SDS–PAGE analysis of 69 designs following IMAC purification. Each sample was run in both unboiled and boiled conditions. Dotted lines indicate the regions corresponding to each backbone group. Positive control (TEV WT) is highlighted in blue, negative control (TEV C151A) in orange, and designs that exhibit cleavage activity ("hits") in red. Samples were analyzed in the same experiment on gels processed in parallel. The experiment was independently repeated 3 times with similar results. **b**, Size-exclusion chromatography (SEC) analysis of 69 designs following IMAC purification. The red dot line indicates the expected elution volume of the cleaved product (free mScarlet). Positive control (TEV WT) is shown in blue box, negative control (TEV C151A) in orange box, and designs that exhibited cleavage activity (“hits”) in red boxes.

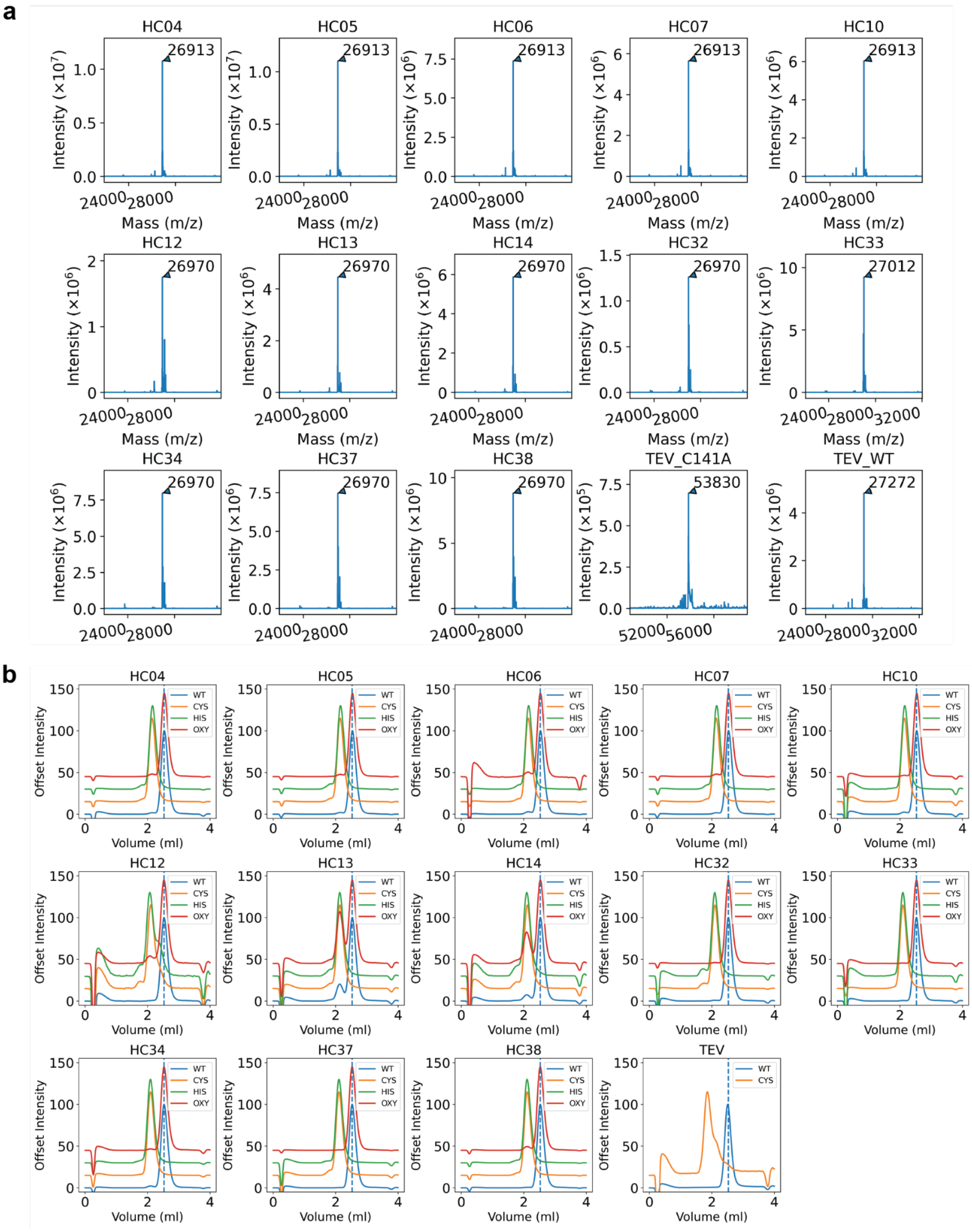

**Extended Data Fig. 3 | LC–MS analysis of cleavage products and knockout analysis of catalytic residues**

**a**, Fractions corresponding to the expected cleaved product were collected from HPLC and analyzed by LC–MS. In all cases, the mass spectra confirmed cleavage at the designed site. **b**, Co-expression screening of designs carrying knockouts of key catalytic residues. Substitution of the cysteine, histidine, or oxyanion hole residue (Asn/Gln) was used to assess intramolecular cleavage activity, confirming their essential roles in catalysis.

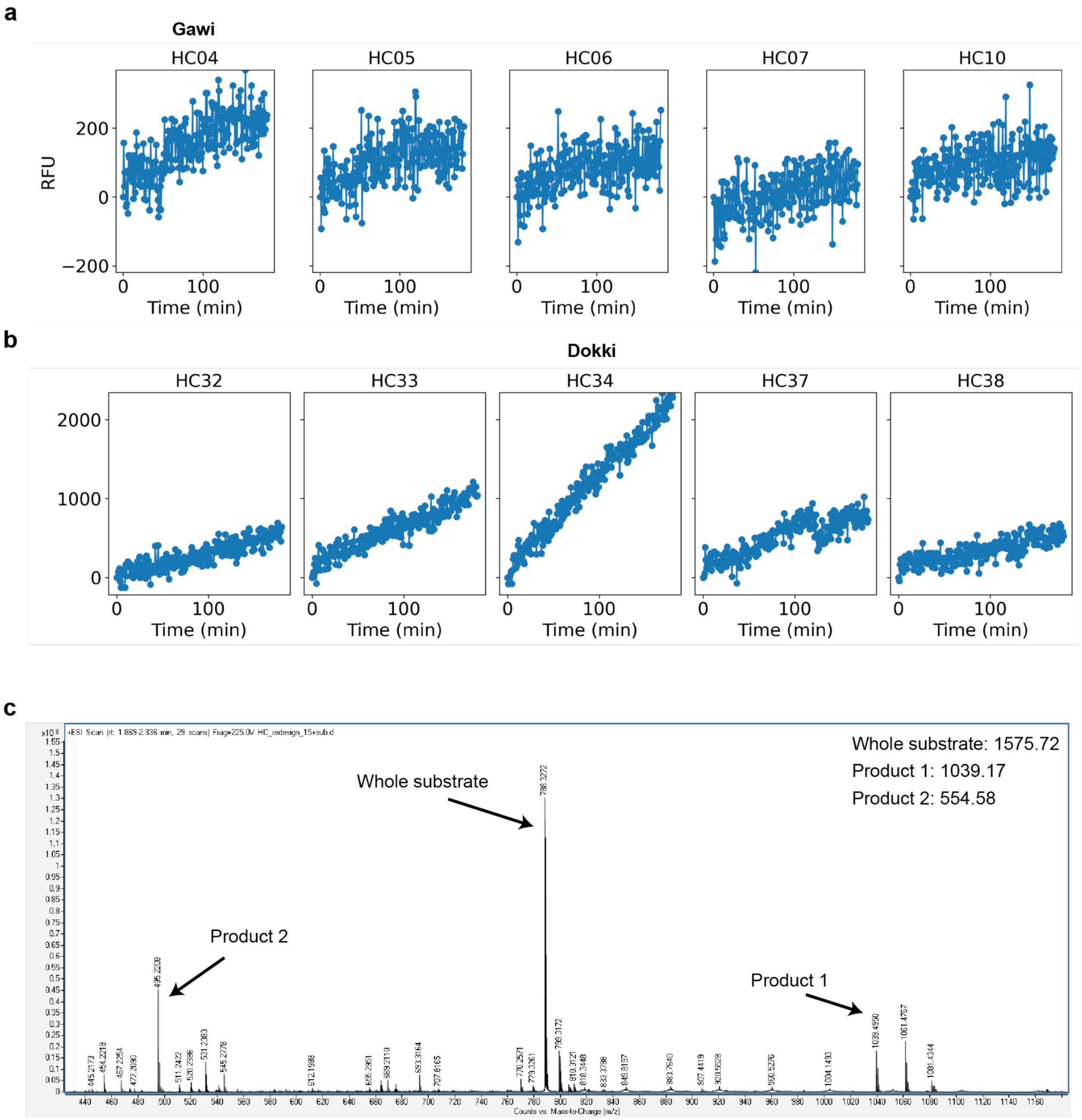

**Extended Data Fig. 4 | Intermolecular cleavage FRET assays of active designs from screening.**

FRET measurements for each design with its cognate FRET-paired peptide substrate. Reactions were performed with 10 µM enzyme and 200 µM substrate in a fluorescence plate reader. **a**, hc04-10. **b**, hc32-38. **c**,LC-MS measurement after FRET assay samples (Dokki) that contain whole substrate, product 1, product 2.

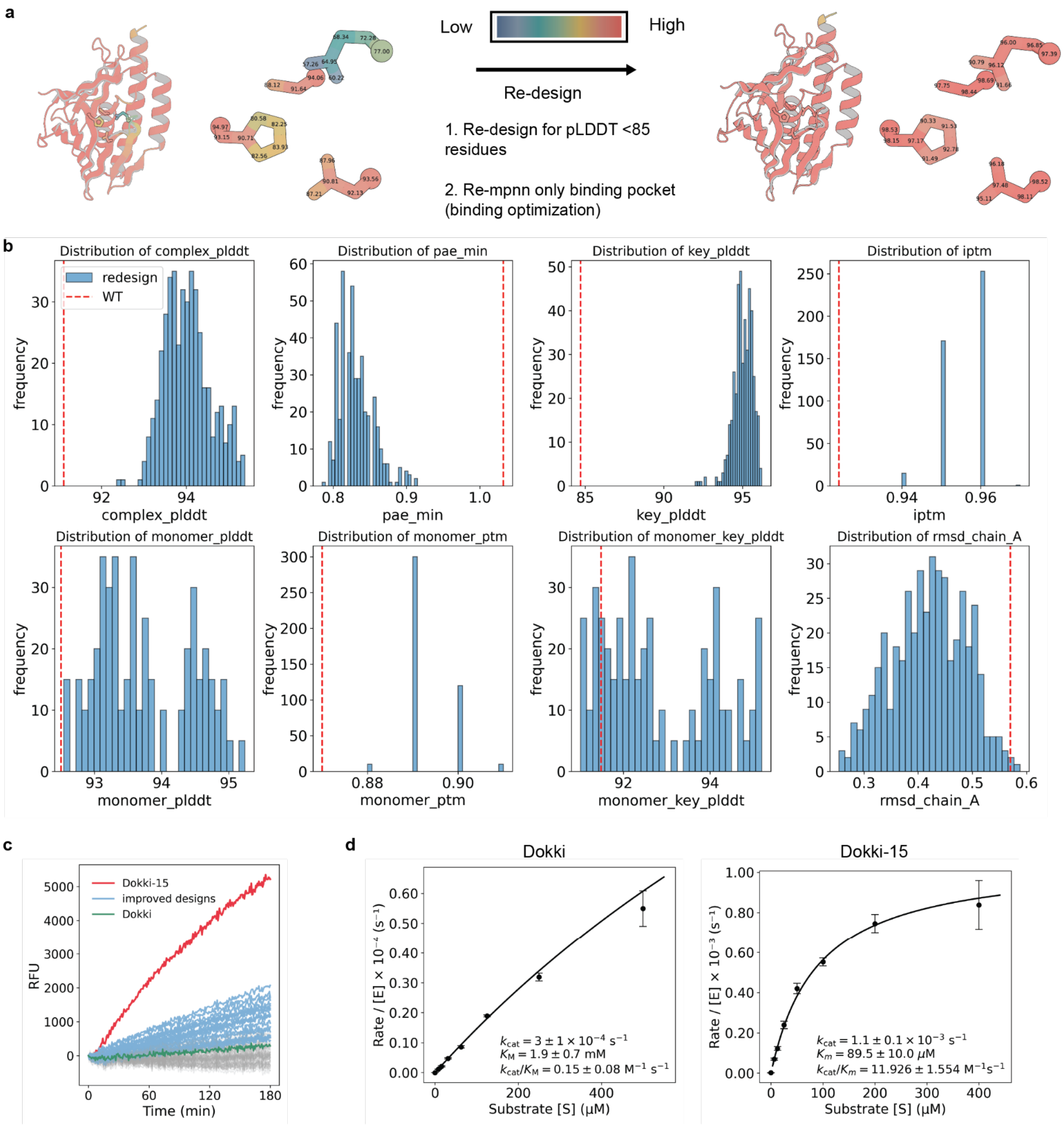

**Extended Data Fig. 5 | Computational re-design of Dokki and catalytic activity comparison to WT.**

**a**, pLDDT-guided re-design workflow applied to Dokki. **b**, Distributions of AF3 prediction metrics for Dokki redesign variants are shown, including complex predictions (pLDDT, min PAE, key-residue pLDDT, iPTM) and apo predictions (pLDDT, pTM, key-residue pLDDT, RMSD to the complex prediction). The red dotted line indicates the AF3 metric values of the

Dokki WT. **c**, Screening reactions were performed with 10 μM enzyme and 100 μM substrate using a plate reader (see Methods). Of 89 redesigns, 24 outperformed the parent Dokki design, with rate improvements ranging from 1.2- to 31-fold. Green indicates WT, blue indicates enhanced designs, and red marks the most improved variant (Dokki-15). **d**, Steady-state kinetic analysis fitted to the Michaelis–Menten equation, providing apparent $k_{cat}$, $K_M$, and catalytic efficiency values of Dokki and Dokki-15. Lines represent the mean and s.d., respectively, of three technical replicates ($n$ = 3).

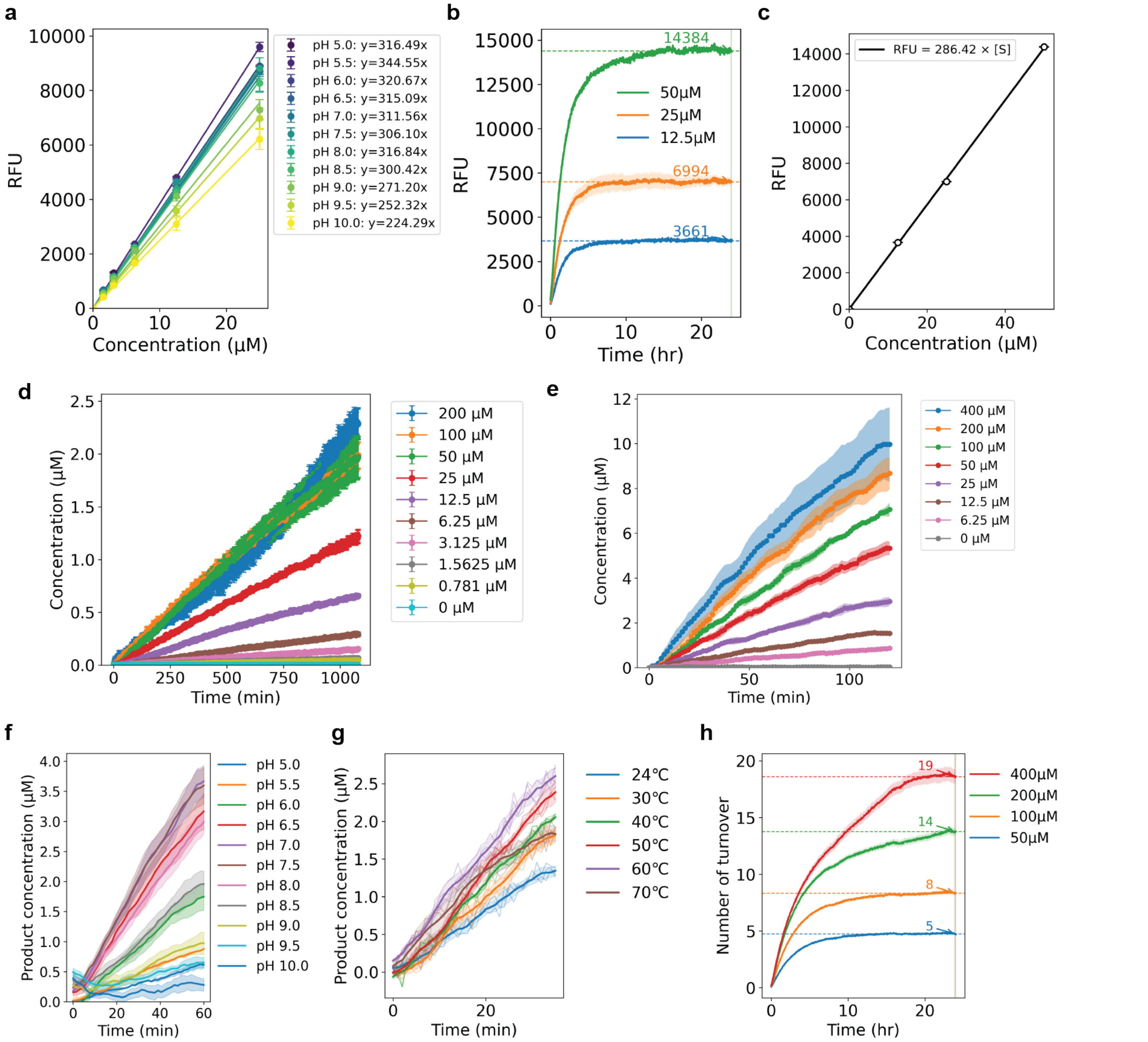

**Extended Data Fig. 6 | Standard curves and characterization of the most active designs.**

**a**, Standard curve for conversion of RFU to free EDANS concentration (50, 25, 12.5, 6.25 μM) (RFU = 311.56 × Conc.(μM)). Lines represent the mean and s.d., respectively, of three technical replicates ($n$ = 3). **b**, Plate reader measurements of complete cleavage products at high enzyme concentration (25 μM) and substrate concentrations of 50, 25, and 12.5 μM, corresponding to

RFU values of 14,384, 6,994, and 3,661, respectively. Lines and shading represent the mean and s.d., respectively, of three technical replicates ($n$ = 3). **c**, Standard curve for conversion of RFU to free GGS–EDANS concentration (50, 25, 12.5 μM) (RFU = 286.42 × Conc.(μM)). Lines represent the mean and s.d., respectively, of three technical replicates ($n$ = 3). **d**, Linear plots for Gawi-09 and **e**, Linear plots for Dokki-15. Lines and shading represent the mean and s.d., respectively, of three technical replicates ($n$ = 3). **f**, Linear plot for pH–rate profiles of Dokki-15. Lines and shading represent the mean and s.d., respectively, of three technical replicates ($n$ = 3). **g**, Linear plots for Dokki-15 as a function of temperature. Lines and shading represent the mean and s.d., respectively, of three technical replicates ($n$ = 3). **h**, Total turnover number (TTN) for different substrate concentrations (400, 200, 100, 50 μM) with 10 μM enzymes, corresponding to maximum TTNs of 40, 20, 10, and 5, respectively. Lines and shading represent the mean and s.d., respectively, of three technical replicates ($n$ = 3).

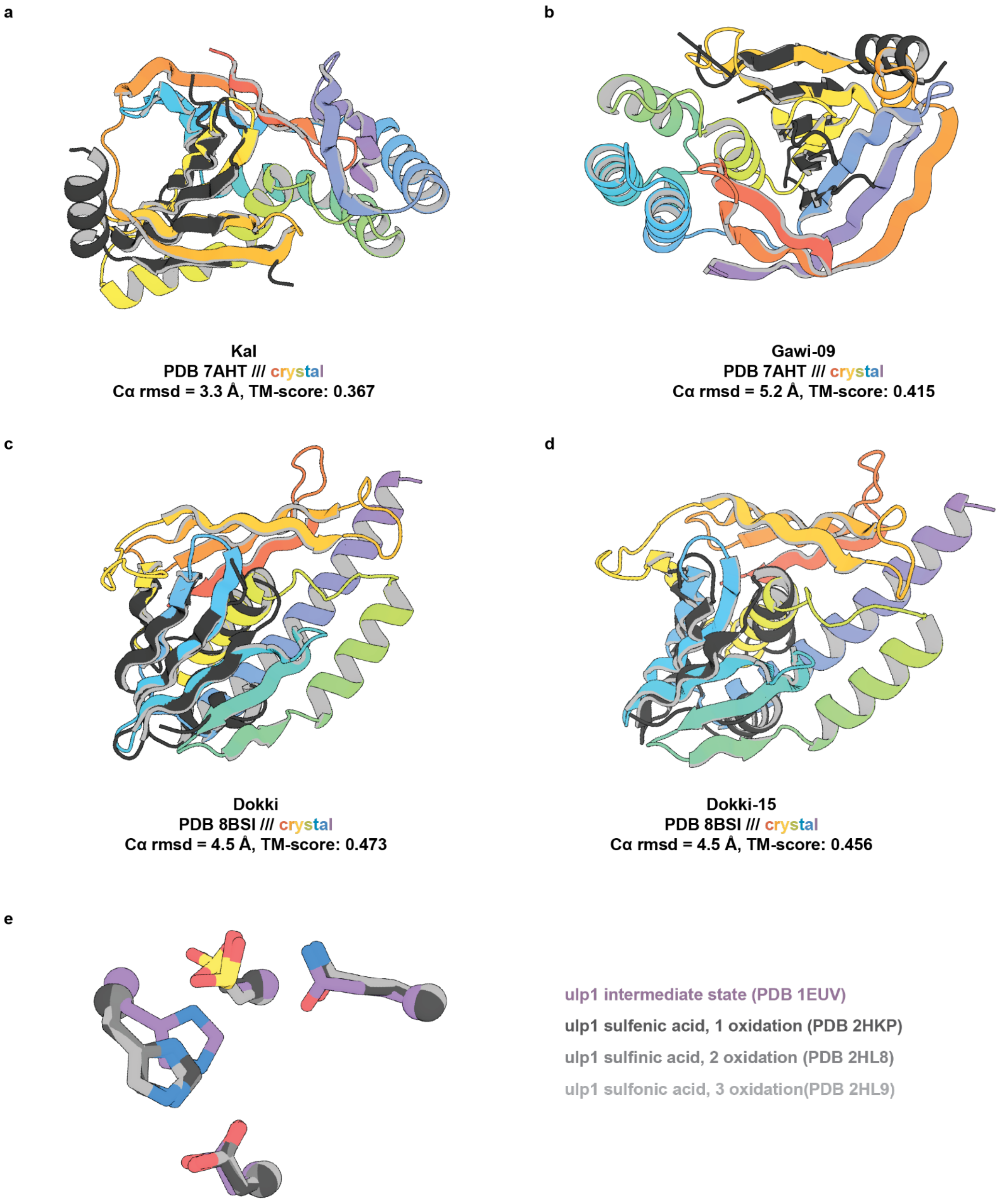

**Extended Data Fig. 7 | Structural similarity analysis of designed cysteine proteases by Foldseek.**

Structural alignment of crystal structures of the designed proteases with their closest matches identified by Foldseek search against the PDB database. For each design, the experimentally determined structure is overlaid with the top-scoring Foldseek hit: **a**, Kal; **b**, Gawi-09; **c**, Dokki; **d**, Dokki-15. Overlays illustrate global fold similarity. These comparisons confirm that the designed scaffolds didn't adopt naturally occurring protein topologies. **e**, Overlay of Ulp1 crystal structures that compare complex intermediate state and oxidation state.

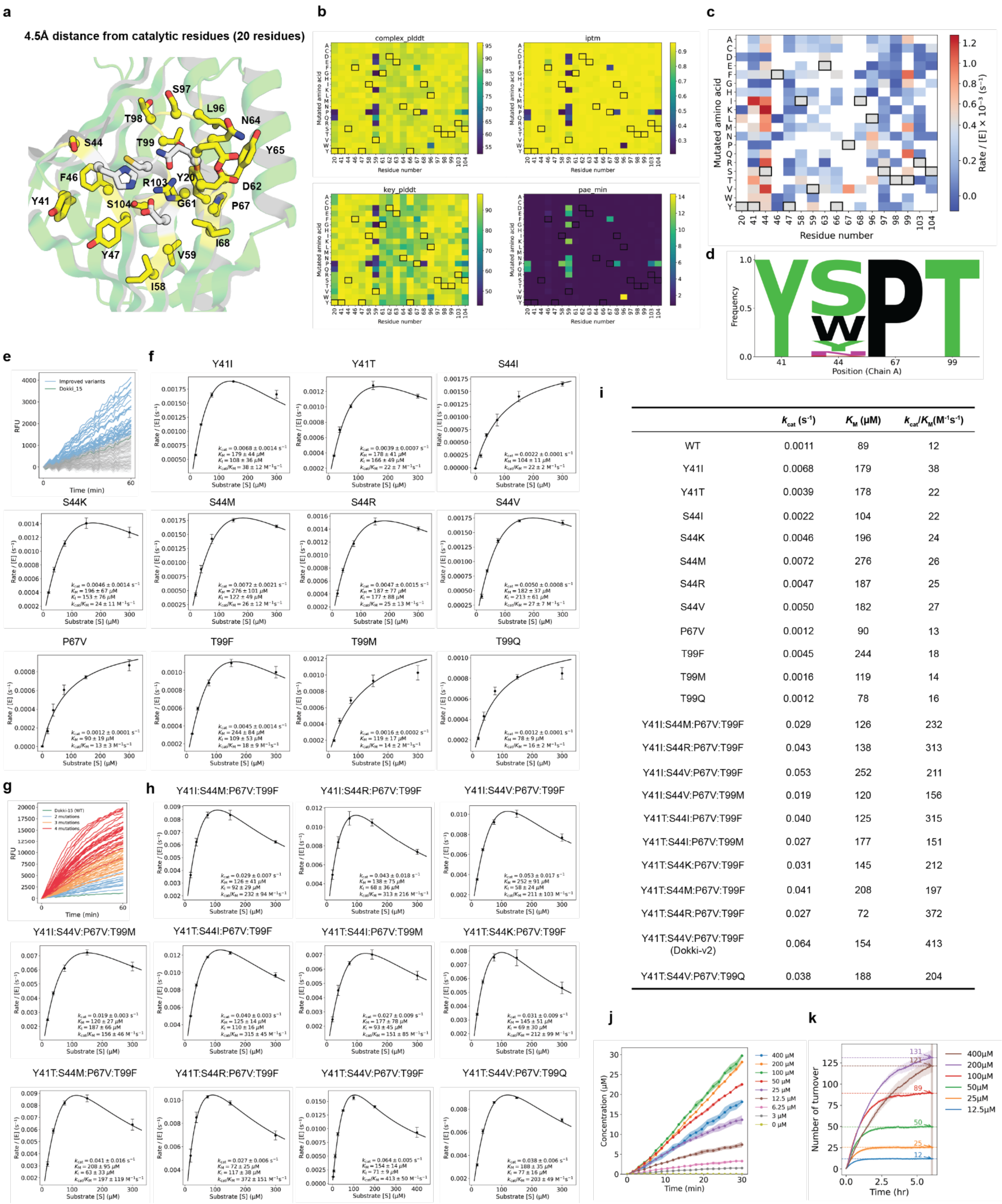

| | $k_{cat}$ ($s^{-1}$) | $K_M$ (μM) | $k_{cat}/K_M$ ($M^{-1}s^{-1}$) |
|---|---|---|---|
| WT | 0.0011 | 89 | 12 |
| Y41I | 0.0068 | 179 | 38 |
| Y41T | 0.0039 | 178 | 22 |
| S44I | 0.0022 | 104 | 22 |
| S44K | 0.0046 | 196 | 24 |
| S44M | 0.0072 | 276 | 26 |
| S44R | 0.0047 | 187 | 25 |
| S44V | 0.0050 | 182 | 27 |
| P67V | 0.0012 | 90 | 13 |
| T99F | 0.0045 | 244 | 18 |
| T99M | 0.0016 | 119 | 14 |
| T99Q | 0.0012 | 78 | 16 |
| Y41I:S44M:P67V:T99F | 0.029 | 126 | 232 |
| Y41I:S44R:P67V:T99F | 0.043 | 138 | 313 |
| Y41I:S44V:P67V:T99F | 0.053 | 252 | 211 |
| Y41I:S44V:P67V:T99M | 0.019 | 120 | 156 |
| Y41T:S44I:P67V:T99F | 0.040 | 125 | 315 |
| Y41T:S44I:P67V:T99M | 0.027 | 177 | 151 |
| Y41T:S44K:P67V:T99F | 0.031 | 145 | 212 |
| Y41T:S44M:P67V:T99F | 0.041 | 208 | 197 |
| Y41T:S44R:P67V:T99F | 0.027 | 72 | 372 |
| Y41T:S44V:P67V:T99F (Dokki-v2) | 0.064 | 154 | 413 |
| Y41T:S44V:P67V:T99Q | 0.038 | 188 | 204 |

**Extended Data Fig. 8 | Mutational screening of Dokki-15 and kinetics for improved variants.**

**a,** Residues selected for mutational screening (yellow) located within 4.5 Å of the catalytic residues (gray). **b,** In silico site saturation mutagenesis evaluated using AF3, showing predicted structural quality metrics for each variant, including complex pLDDT, ipTM, key-residue pLDDT, and minimum predicted aligned error (PAE). **c,** Heat map summarizing wet-lab mutational screening. **d,** Sequence logo representing amino acid frequencies at positions corresponding to beneficial mutations in ProteinMPNN-generated sequences. **e,** FRET-based activity screen of single-mutation variants measured with 3 μM enzyme and 75 μM substrate over 1 h. Dokki-15 is shown in green, and activity-enhancing variants are shown in blue. **f,** Michaelis–Menten plots for selected beneficial single-mutation variants. Lines represent the mean and s.d., respectively, of three technical replicates (*n* = 3). **g,** FRET-based activity screen of combinatorial variants measured with 3 μM enzyme and 75 μM substrate over 1 h. Variants containing two mutations are shown in blue, three mutations in orange, and four mutations in red. **h,** Michaelis–Menten plots for selected combinatorial variants. Lines represent the mean and s.d., respectively, of three technical replicates (*n* = 3). **i,**Summary of catalytic parameters ($k_{cat}$, $K_M$, and $k_{cat}/K_M$) for mutants. **j,** Linear plots for Dokki-v2. Lines and shading represent the mean and s.d., respectively, of three technical replicates (*n* = 3). **k**, Total turnover number (TTN) for different substrate concentrations (400, 200, 100, 50, 25, 12.5 μM) with 1 μM enzymes, corresponding to maximum TTNs of 400, 200, 100, 50, 25 and 12.5, respectively. Lines and shading represent the mean and s.d., respectively, of three technical replicates (*n* = 3).

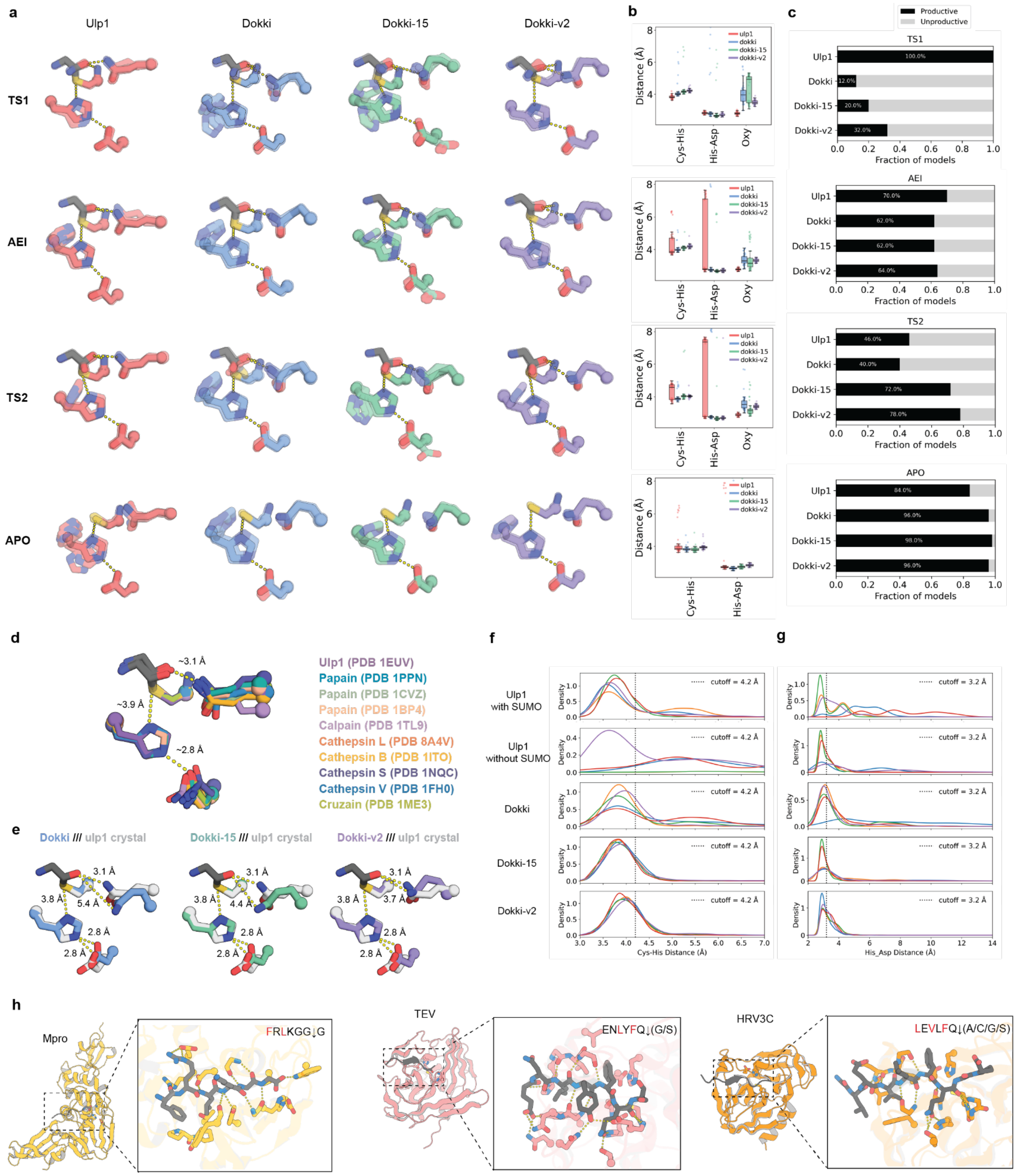

**Extended Data Fig. 9 | Computational analysis of dokki-series designs compared to native cysteine proteases.**

**a,** Representative PLACER ensembles of AEI, TS2 and apo states, showing the catalytic triad and the oxyanion-hole residue Gln. **b,** Distributions of Cys–His, His–Asp and oxyanion-hole distances across PLACER ensembles for Ulp1 and the Dokki series in the indicated states ($n$ = 50 models generated using 50 independent random seeds per design and state). Boxes show the median and 25th–75th percentiles; whiskers extend to the most extreme values within 1.5 × the interquartile range, and points represent individual models. **c,** Fraction of productive conformations in TS1-, AEI-, TS2- and apo-state PLACER ensembles. **d,** Overlay of catalytic-site structures from representative native papain-like cysteine proteases. **e,** Comparison of active-site geometries in the Ulp1 crystal structure and AF3 models of Dokki, Dokki-15 and Dokki-v2, showing stepwise improvement in oxyanion-hole geometry. **f,** Density distributions from molecular dynamics simulations showing distances between the catalytic cysteine sulfur atom and His ND1, and **g,** His NE2 and Asp OD1. Each system was simulated in five independent 500-ns replicas, shown as separate traces. **h,** Substrate-binding pockets of Mpro, TEV and HRV3C, with residues forming hydrophobic pockets highlighted in red.

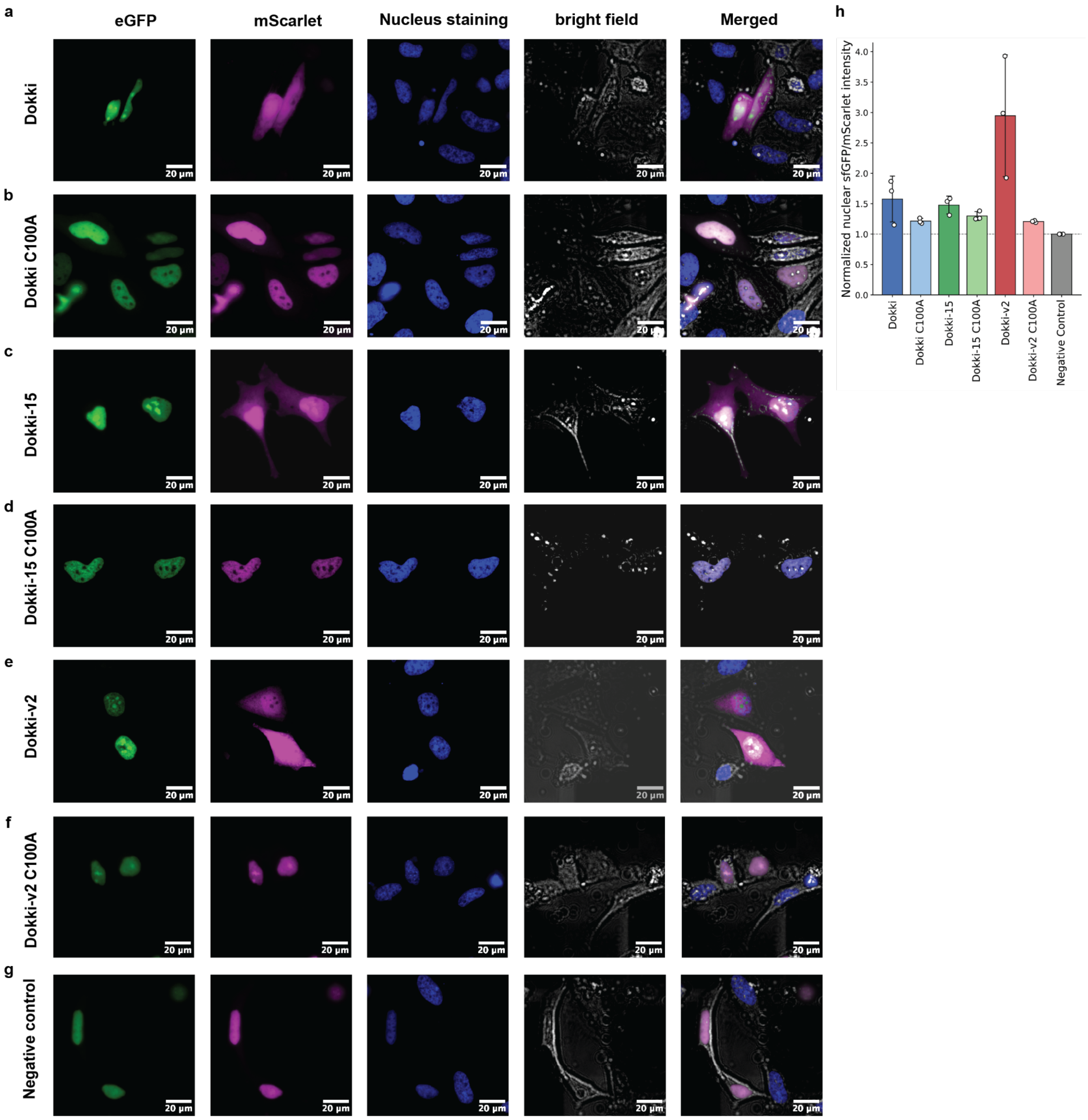

**Extended Data Fig. 10 | Cell-based reporter assay for enzyme-dependent substrate relocalization.**

**a–g,** Representative fluorescence microscopy images of HeLa cells showing sfGFP, mScarlet, Nucleus staining, bright field and merged channels 24 h after transfection. Substrate and protease plasmids were transfected at a 100 ng:10 ng ratio before imaging by InCell microscopy. **a,** Dokki; **b,** Dokki C100A; **c,** Dokki-15; **d,** Dokki-15 C100A; **e,** Dokki-v2; **f,** Dokki-v2 C100A; **g,** Negative control. **h,** Quantitative data for the sfGFP/mScarlet signal inside the nucleus that is normalized by negative control. **h,** Nuclear sfGFP/mScarlet fluorescence ratios normalized to the negative-control condition. Each dot represents one biologically independent experiment; bars show the mean and error bars indicate s.d. across the three experiments ($n$ = 3). On average 200 cells per condition were included across the three experiments. C100A denotes the catalytically inactive active-site mutant.

### Supplementary Information Guide

**Supplementary Table 1. Sequences of designed proteases and substrates**
Amino acid sequences of the designed proteases and their corresponding substrate sequences, with cleavage sites indicated.

**Supplementary Table 2. LC–MS data**
Expected and observed molecular masses of substrate cleavage products, and their mass differences, for the tested protease designs.

**Supplementary Table 3. Crystallography data collection and refinement statistics**
X-ray diffraction data collection and structure refinement statistics for six apo and substrate-bound crystal structures of the designed proteases.

**Supplementary Table 4. Foldseek analysis of crystal structures**
Structural matches identified by Foldseek, including TM-scores, source databases and target identifiers for the designed protease crystal structures.

**Supplementary Table 5. Kinetic summary of sequence-specific cysteine proteases**
Substrates and kinetic parameters ($k_{cat}$, $K_M$ and $k_{cat}$ / $K_M$) for designed and reference cysteine proteases.

**Supplementary Figure 1. Full SDS–PAGE gel images**
Full gel images corresponding to Fig. 1d and Extended Data Fig. 2.

**Supplementary Figure 2. Gel-based kinetic analysis of Ulp1 cleavage activity**
Gel-based cleavage assays and kinetic analysis of Ulp1 acting on eGFP–peptide–mScarlet substrates with or without SUMO, with three technical replicates per substrate concentration.

**Supplementary Figure 3. Uncropped images for the cell-based reporter assay**
Representative uncropped fluorescence microscopy images of HeLa cells for Dokki, Dokki-15, Dokki-v2, their C100A mutants and the negative control. White boxes indicate the cropped regions.

**Supplementary Information 1. pCool6 plasmid DNA sequence**
Nucleotide sequence of the pCool6 plasmid used for initial screening in *Escherichia coli*. The plasmid encodes an insert–mScarlet–His-tag fusion under the control of a T7 promoter and contains a kanamycin resistance gene.

**Supplementary Information 2. ZJ101 plasmid DNA sequence**
Nucleotide sequence of the ZJ101 plasmid used for mammalian cell expression. The plasmid contains a CMV promoter, a γ-globin intron and an ampicillin resistance gene.

#### Supplementary table 1. Sequences for designed protease and substrate

| design_id | design_aa_seq | substrate sequence |
|---|---|---|
| **HC04 (Gawi)** | TVKSFPATVTEYTTADGKTYYEIKLKDFSLTDEELEKIYQKVAETDPENKEKITAYGKALAEAG<br>VASKADPEKLVALAKKYGVTDRVCLLVALLYIQADPSWSLFYTPNNHFVALKVDADGKVLVR<br>DLNSNNTALTESSYPSREAFLASLGIVSSKTFETTVSLTASNGLPAIKIE | GSKIISGG/S |
| **HC05** | TVKSFPATVTEYTTADGKTYYEIKLKDFSLTDEELEKIYQKVAETDPENKEKITAYGKALAEAG<br>VASKADPEKLVALAKKYGVTDRVCLLVALLYIQADPSWSLFYTPNNHFVALKVDADGKVLVR<br>DLNSNQTALTESSYPSREAFLASLGIVSSKTFETTVSLTASNGLPAIKIE | GSKIISGG/S |
| **HC06** | TVKSFPATVTKYTTADGKEYYEIDLKDFKLTDEELAAIYDKVAETDPSRKEAITAYGKALAEA<br>GVASKADPEKLVALAKQYGVTDRICLLVALLYIQADPSWKLFYTPNNHFVAIKEDENGKVLVK<br>DLNANNTALTERSYPSREAFLKSLGIVSSETFETTVSLTASNGLPAIKIE | GSKIISGG/S |
| **HC07** | TVKSFPATVTKYTTADGKEYYEIDLKDFKLTDEELAAIYDKVAETDPSRKEAITAYGKALAEA<br>GVASKADPEKLVALAKQYGVTDRICLLVALLYIQADPSWKLFYTPNNHFVAIKEDENGKVLVK<br>DLNANQTALTERSYPSREAFLKSLGIVSSETFETTVSLTASNGLPAIKIE | GSKIISGG/S |
| **HC10** | VVKSFPATVTKYTTADGKEYYEIDLKDFRLTDEELKAIYDKVAETDPSRADAIRAYGDALAEA<br>KVAEKADPELLVALAKEYGVTDIVCLLVALLYIQADPSWKLFYTPNNHFVALKVDENGKVTV<br>RDLNSNNTALTERSYPSREAFLSSLGIVSSETFETTVSLTASNGLPAIKIE | GSKIISGG/S |
| **HC12 (Kal)** | ATQTLTVKVAPGVEVEVSFEGTEEQREKAQAIVDEGNKTLDSVDLGKGKLYKDKNGNIIVEISR<br>EQLGKAAELAKTRLPYIDGVAAAGAPGGSCLPLSIIRAYEEYYKAEELGKELGVPVSDKVLVV<br>NFNNQHFVVQVPVGDKLYVFDVDENNNPVYYTEPLPELTYVADVNVTFYASGISGSIEAIP | SIVLTGSG/S |
| **HC13** | ATQTLTVEVAPGVSVEVKFEGTEEQREKAQAIIDEAQKNLSSVNLGKGKLYKDKEGNTLVEIT<br>VEQLKKAAELAKTIKPAIDKVKALGAAGGSCLPLGVIRAVERYEKLKEEAKKKGLEVSETVFIK<br>NINDSHFVILIPVGDKWYVVDFNEKNEPVEYFIDKPELTYVGDVDVTFWVGGISGAIEITP | SIVLTGSG/S |
| **HC14** | ATQTLTVRVAPGVSVTVEFEGTEAQRDAAQAIIAEGEKILSSVDLGKGRLYRDKEGNIVVEITR<br>EQLEKAAELAKKLLPYIDAVAAAGAPGGSCLPLSLIEAVKKIEELEEEAKKKGLEVSEKVYIVNI<br>DNSHFVVLIPVGDKAFVVDFDANNNPVLSEVPLPELEFVGEVNITFYAGGISGAIKADL | SIVLTGSG/S |
| **HC32** | SLEERYKKRAEGVLEIYEVYKEVGKELAKKYPDAEVQLVNYNNSHFYLRLNTSKGVVTVDGD<br>ENNNPVVTDAGKTTYDEKAKEKLDKIAEEYDIDLSTTCLIRSLAIIEAFYEVNDGKSPVSLVTEG<br>DGVPLQEVSDANLSGKVYKKNDKSEYLELKGSWNSDGKVTLEIRNPKGEVLAKIDIKV | VPTLTGG/G |
| **HC33** | SEKAEYEKRAEGVKEIYEVYYETGKKLAEEYPDATVNLVNYDNSHFYLELETPEGTVIVDGDE<br>NNYPVVTKDGKKTVDEEAKEKLKEIAEKYDIDLSTTCLIRSLAIVQAYWETRNGKAPVRLYTE<br>GTGVPRQEVAEANLSGKIYKKDDKSKYLKVTGAWRPDGVLTLNLLDPEGNVLASYTIRY | VPTLTGG/G |
| **HC34 (Dokki)** | SEKAEYEKRAEGVKEIYEVYYETGKKLAEEYPDATVNLVNYDNSHFYLELETPEGTVIVDGDE<br>NQYPVVTKDGKKTVDEEAKEKLKEIAEKYDIDLSTTCLIRSLAIVQAYWETRNGKAPVRLYTE<br>GTGVPRQEVAEANLSGKIYKKDDKSKYLKVTGAWRPDGVLTLNLLDPEGNVLASYTIRY | VPTLTGG/G |
| **HC37** | SAKEEQKKRVEGAVEIYKVYYEAGKKLAKKYPDAKVKLVNYDNSHFYLELETPKGTVKVDG<br>DSNNNIVVTEGGKKTVDEEAKKKLDEIAEKYNIDLSTTCLIRSLAIIKAYQETNNGKTPVELYTE<br>GTGVPKKKIDEANLSGKIKEKNDSSKYINLKGSWNKDGVITLELLDENGEVLKKIEIKV | VPTLTGG/G |
| **HC38** | SAKEEQKKRVEGAVEIYKVYYEAGKKLAKKYPDAKVKLVNYDNSHFYLELETPKGTVKVDG<br>DSNQNIVVTEGGKKTVDEEAKKKLDEIAEKYNIDLSTTCLIRSLAIIKAYQETNNGKTPVELYTE<br>GTGVPKKKIDEANLSGKIKEKNDSSKYINLKGSWNKDGVITLELLDENGEVLKKIEIKV | VPTLTGG/G |
| **Gawi-09** | TVKSFPATVTKYTTADGKEYYEVDLKDFKLTDEELAAIYDKVAETDPSRKEAITAYGKALAEA<br>GVASKADPEKLVALAKQYGVTDRICLLVALLYIQADPSWKVFFTPNNHAVAIKEDENGKVLV<br>KDLNANNTALTERSYPSREAFLKSLGIVSSETFESTASLTASDGLPAIEVD | VPTLTGG/G |
| **Dokki-15** | GLDAEYEKLAEGVKEIYEVYYETGKKLAEEYPDATVNLVNYDNSHFYLELETPKGTVIVDGDE<br>NQYPIVTKNGKKTEDKEAKAKLKEIAEKYDIDLSTTCLIRSLAIVQAYWETRNGKAPVKLYSEG<br>TGVPRQELAAANLVSKIYKKGDKSKYLKVTGAWRPDGVLTLNLLDPNGNVLASYTIKY | VPTLTGG/G |
| **Dokki-v2** | GLDAEYEKLAEGVKEIYEVYYETGKKLAEEYPDATVNLVNTDNVHFYLELETPKGTVIVDGDE<br>NQYVIVTKNGKKTEDKEAKAKLKEIAEKYDIDLSTFCLIRSLAIVQAYWETRNGKAPVKLYSEG<br>TGVPRQELAAANLVSKIYKKGDKSKYLKVTGAWRPDGVLTLNLLDPNGNVLASYTIKY | VPTLTGG/G |

**Supplementary table 2. LC-MS data**

| Designs | Expected cleaved MW (Da) | Observed MW (Da) | Difference (Da) |
|---|---|---|---|
| **hc04 (Gawi)** | 26913.33 | 26913.06 | 0.27 |
| **hc05** | 26913.33 | 26913.07 | 0.26 |
| **hc06** | 26913.33 | 26913.05 | 0.28 |
| **hc07** | 26913.33 | 26913.06 | 0.27 |
| **hc10** | 26913.33 | 26913.06 | 0.27 |
| **hc12 (Kal)** | 26970.38 | 26969.99 | 0.39 |
| **hc13** | 26970.38 | 26970 | 0.38 |
| **hc14** | 26970.38 | 26970.03 | 0.35 |
| **hc32** | 26970.38 | 26970.02 | 0.36 |
| **hc33** | 26970.38 | 26970.4 | -0.02 |
| **hc34 (Dokki)** | 26970.38 | 26970.04 | 0.34 |
| **hc37** | 26970.38 | 26970.05 | 0.33 |
| **hc38** | 26970.38 | 26970.04 | 0.34 |
| **TEV** | 27272.67 | 27272.17 | 0.5 |
| **TEV C141A** | 53830.1849 | 53830.22 | -0.0351 |

**Supplementary table 3. Crystallography data collection and refinement statistics**

| | **Kal WT complex**<br>PDB Code: 9YOZ | **Kal C103A complex**<br>PDB Code: 9YOY | **Dokki WT apo**<br>PDB Code: 9YP0 |
|---|---|---|---|
| **Data collection** | | | |
| Space group | $P\ 4_3\ 2_1\ 2$ | $P\ 4_3\ 2_1\ 2_1$ | $P\ 2_1\ 2_1\ 2_1$ |
| Cell dimensions | | | |
| *a*, *b*, *c* (Å) | 77.23, 77.23, 104.97 | 77.29, 77.29, 104.66 | 37.44, 57.04, 75.75 |
| α, β, γ (˚) | 90, 90, 90 | 90, 90, 90 | 90, 90, 90 |
| Resolution (Å) | 104.97 - 2.19 (2.26 - 2.19) | 104.66 - 1.96 (2.01 - 1.96) | 37.88 - 1.77 (1.86 - 1.77) |
| $R_{sym}$ or $R_{merge}$ | 0.437 (1.518) | 0.105 (1.956) | 0.284 (1.875) |
| $I / \sigma I$ | 11.3 (2.7) | 21.1 (2.0) | 9.5 (2.0) |
| Completeness (%) | 100 (100) | 100 (100) | 99.9 (99.6) |
| Redundancy | 25.9 (27.6) | 25.9 (26.6) | 9.9 (10.2) |
| **Refinement** | | | |
| Resolution (Å) | 62.21 - 2.19 (2.33 - 2.19) | 62.17 - 1.96 (2.05 - 1.96) | 37.88 - 1.77 (1.82 - 1.77) |
| No. reflections | 16675 (2734) | 22604 (2699) | 0.2452 (0.3386) |
| $R_{work}$ / $R_{free}$ | 0.1976 (0.3072) / 0.2546 (0.3361) | 0.1857 (0.3229) / 0.2193 (0.3577) | 0.1973 (0.3160)/ 0.2452 (0.3386) |
| No. atoms | | | |
| Protein | 1517 | 1528 | 1507 |
| Ligand/ion | n/a | n/a | n/a |
| Water | 69 | 64 | 54 |
| *B*-factors | | | |
| Protein | 47 | 55 | 21 |
| Ligand/ion | n/a | n/a | n/a |
| Water | 46 | 52 | 29 |
| R.m.s. deviations | | | |
| Bond lengths (Å) | 0.005 | 0.013 | 0.012 |
| Bond angles (°) | 0.71 | 1.22 | 1.15 |

*Single xtal used for each data/structure.
*Values in parentheses are for highest-resolution shell.

Cont…

| | **Gawi-09 WT apo**<br>PDB Code: 9YOX | **Dokki-15 WT apo**<br>PDB Code: 9YNM | **Dokki-15 C100A complex**<br>PDB Code: 9YNL |
|---|---|---|---|
| **Data collection** | | | |
| Space group | P 1 $2_1$ 1 | P $2_1$ $2_1$ $2_1$ | P $4_1$ $2_1$ 2 |
| Cell dimensions | | | |
| *a*, *b*, *c* (Å) | 42.28, 61.19, 63.36 | 38.23, 61.86, 74.76 | 61.63, 61.63, 374.44 |
| α, β, γ (˚) | 90, 105.24, 90 | 90, 90, 90 | 90, 90, 90 |
| Resolution (Å) | 32.88 - 2.17 (2.28 - 2.17) | 34.04 - 1.34 (1.42 - 1.34) | 34.48 - 3.38 (3.56 - 3.38) |
| $R_{sym}$ or $R_{merge}$ | 0.242 (2.212) | 0.094 (1.888) | 0.755 (4.883) |
| $I$ / σ$I$ | 5.7 (1.6) | 11.0 (0.4) | 5.0 (1.4) |
| Completeness (%) | 99.5 (98.8) | 85.6 (46.1) | 99.9 (100) |
| Redundancy | 7.0 (7.0) | 11.4 (4.7) | 25.2 (27.0) |
| **Refinement** | | | |
| Resolution (Å) | 32.88 - 2.17 (2.23 - 2.17) | 34.04 - 1.56 (1.61 - 1.56) | 34.48 - 3.38 (3.53 - 3.38) |
| No. reflections | 16577 (1386) | 25703 (2197) | 10991 (1315) |
| $R_{work}$ / $R_{free}$ | 0.2079 (0.2886) / 0.2655 (0.3424) | 0.1715 (0.2117) / 0.2017 (0.2625) | 0.2226 (0.2974) / 0.2794 (0.3333) |
| No. atoms | | | |
| Protein | 2868 | 1512 | 4543 |
| Ligand/ion | 10 | n/a | n/a |
| Water | 55 | 207 | 6 |
| *B*-factors | | | |
| Protein | 38 | 22 | 94 |
| Ligand/ion | 47 | n/a | n/a |
| Water | 37 | 34 | 56 |
| R.m.s. deviations | | | |
| Bond lengths (Å) | 0.002 | 0.010 | 0.002 |
| Bond angles (°) | 0.48 | 1.15 | 0.44 |

*Single xtal used for each data/structure.
*Values in parentheses are for highest-resolution shell.

**Supplementary table 4. Foldseek analysis for crystal structures**

| Designs | TM score | Database | Target |
|---|---|---|---|
| **Kal WT complex** | 0.48 | AFDB50 | AF-A0A923ZJ62-F1-model_v6 |
| **Kal C103A complex** | 0.49 | AFDB50 | AF-A0A923ZJ62-F1-model_v6 |
| **Gawi-09 WT apo** | 0.46 | AFDB50 | AF-A0A0X8G3I4-F1-model_v6 |
| **Dokki WT apo** | 0.50 | All databases | A0A346A683 |
| **Dokki-15 WT apo** | 0.50 | MGNIFY_ESM30 | MGYP001077622663 |
| **Dokki-15 C100A complex** | 0.48 | CATH50 | 2.170.210.20 Q9N4L1 1-108 |

**Supplementary table 5. Kinetics summary of sequence specific cysteine proteases**

| Enzymes | substrate | $k_{cat}$ ($s^{-1}$) | $K_M$ (μM) | $k_{cat}/K_M$ ($M^{-1}$ $s^{-1}$) |
|---|---|---|---|---|
| **Dokki** | DABCYL–GSVPTLTGG↓GS – EDANS | 0.0003 ± 0.0001 | 1900 ± 700 | 0.15 ± 0.08 |
| **Dokki-15** | DABCYL–GSVPTLTGG↓GS – EDANS | 0.0011 ± 0.0001 | 89 ± 10 | 12 ± 2 |
| **Dokki-v2** | DABCYL–GSVPTLTGG↓GS – EDANS | 0.064 ± 0.005 | 154 ± 14 | 413 ± 50 |
| **Ulp1** | eGFP– SUMO-REQIGG↓G–mScarlet | 6.1 ± 0.5 | 17 ± 2 | $3.5 \pm 0.4 \times 10^5$ |
| | eGFP–REQIGG↓G–mScarlet | Not measurable | Not measurable | Not measurable |
| **TEV(Kapust et al. 2001)** | ENLYFQ↓S | 0.16 ± 0.01 | 61 ± 10 | 2620 ± 460 |
| **mPro(Kenneson et al. 2025)** | DABCYL–KTSAVLQ↓SGFRKME–(EDANS)-NH2 | 1.23 | 190 | 6,600 |
| **HRV3C(Kenneson et al. 2025)** | MBP–LEVLFQ↓X-6His–GFP/GST | 0.021-0.32 | 34 - 49 | 390 - 4730 |

**a. Figure 1d: hc04,05,06,07,10,12,13,14,32,33,34,37,38,TEV WT, TEV C151A**

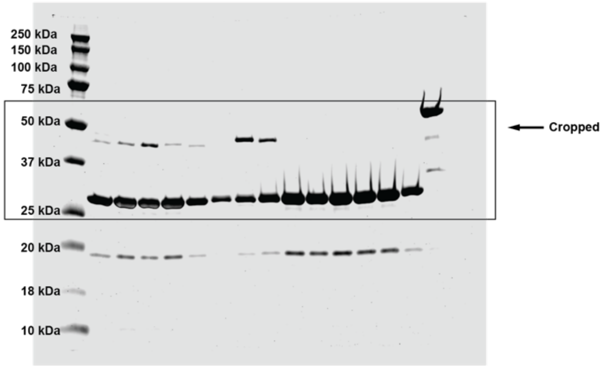

**b. Extended data figure 2: hc01-12**

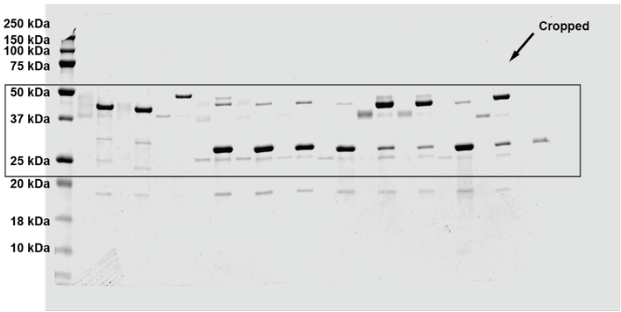

**c. Extended data figure 2: hc13-24**

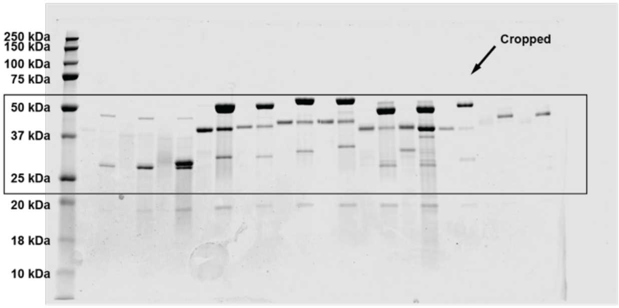

**d. Extended data figure 2: hc25-36**

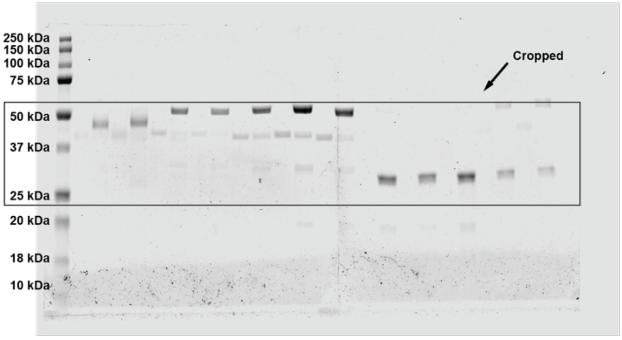

**e. Extended data figure 2: hc37-48**

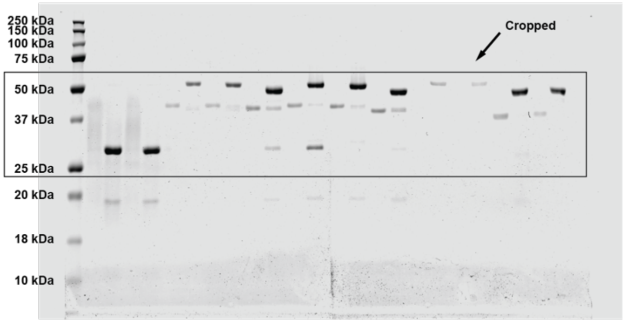

**f. Extended data figure 2: hc49-60**

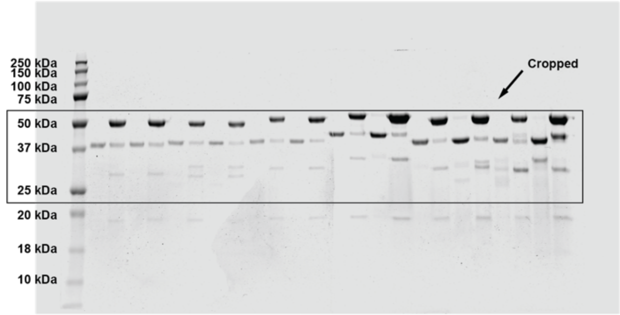

**g. Extended data figure 2: hc61-69 & TEV WT, TEV C151A**

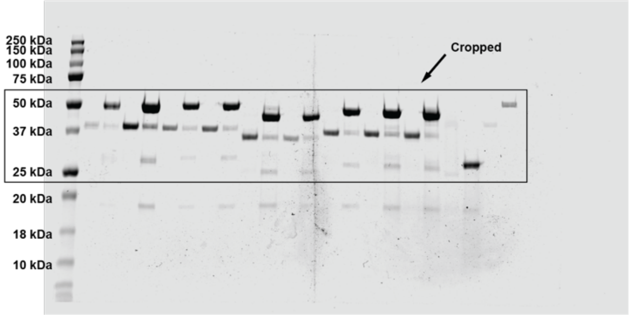

**Supplementary Fig. 1 | Full image for SDS-page gel.**

Full SDS-page gel image for **a,** Figure 1d, **b–g,** Extended data Figure 2

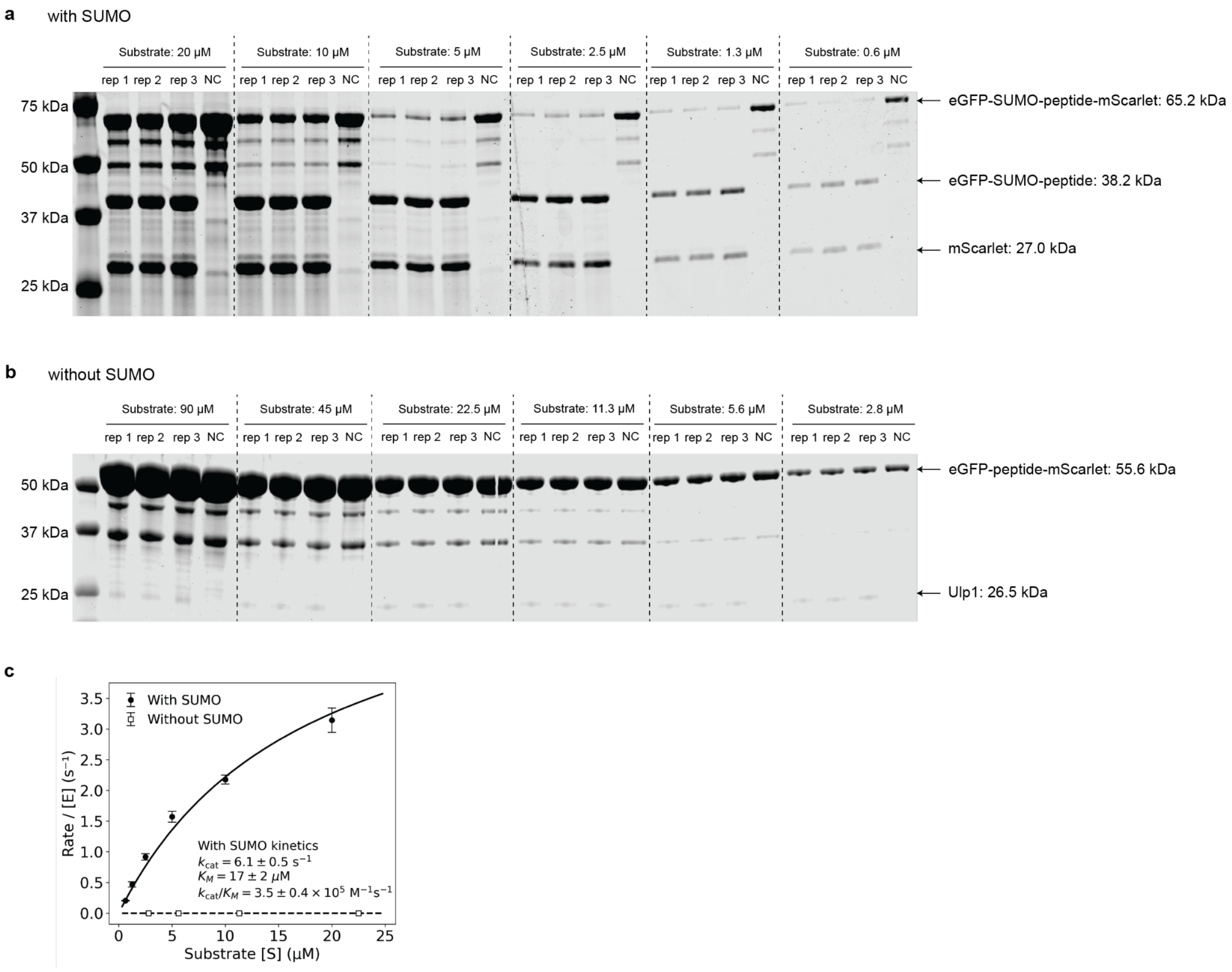

**Supplementary Fig. 2 | Gel-based kinetic analysis of Ulp1 cleavage activity.**

Ulp1 activity against eGFP–peptide–mScarlet substrates was measured using a gel-based cleavage assay. **a,** For SUMO-containing substrates, reactions contained 30 nM Ulp1 and two-fold serial dilutions of substrate from 20 to 0.6 μM. Reactions were incubated for 1 min at room temperature. Each concentration performed three technical replicates. **b,** For substrates lacking SUMO, reactions contained 1 μM Ulp1 and two-fold serial dilutions of substrate from 90 to 2.8 μM. Reactions were incubated for 1 h at room temperature. Each concentration performed three technical replicates. Reaction products were resolved by SDS–PAGE, stained with Coomassie blue and imaged on a LI-COR imaging system using Coomassie detection. **c,** Steady-state kinetic analysis of the Ulp1 to with/without SUMO substrate, fitted to the Michaelis–Menten substrate inhibition model to derive apparent $k_{cat}$, $K_M$, $K_I$, and catalytic efficiency ($k_{cat}/K_M$). Error bars represent the standard deviation of three technical replicates.

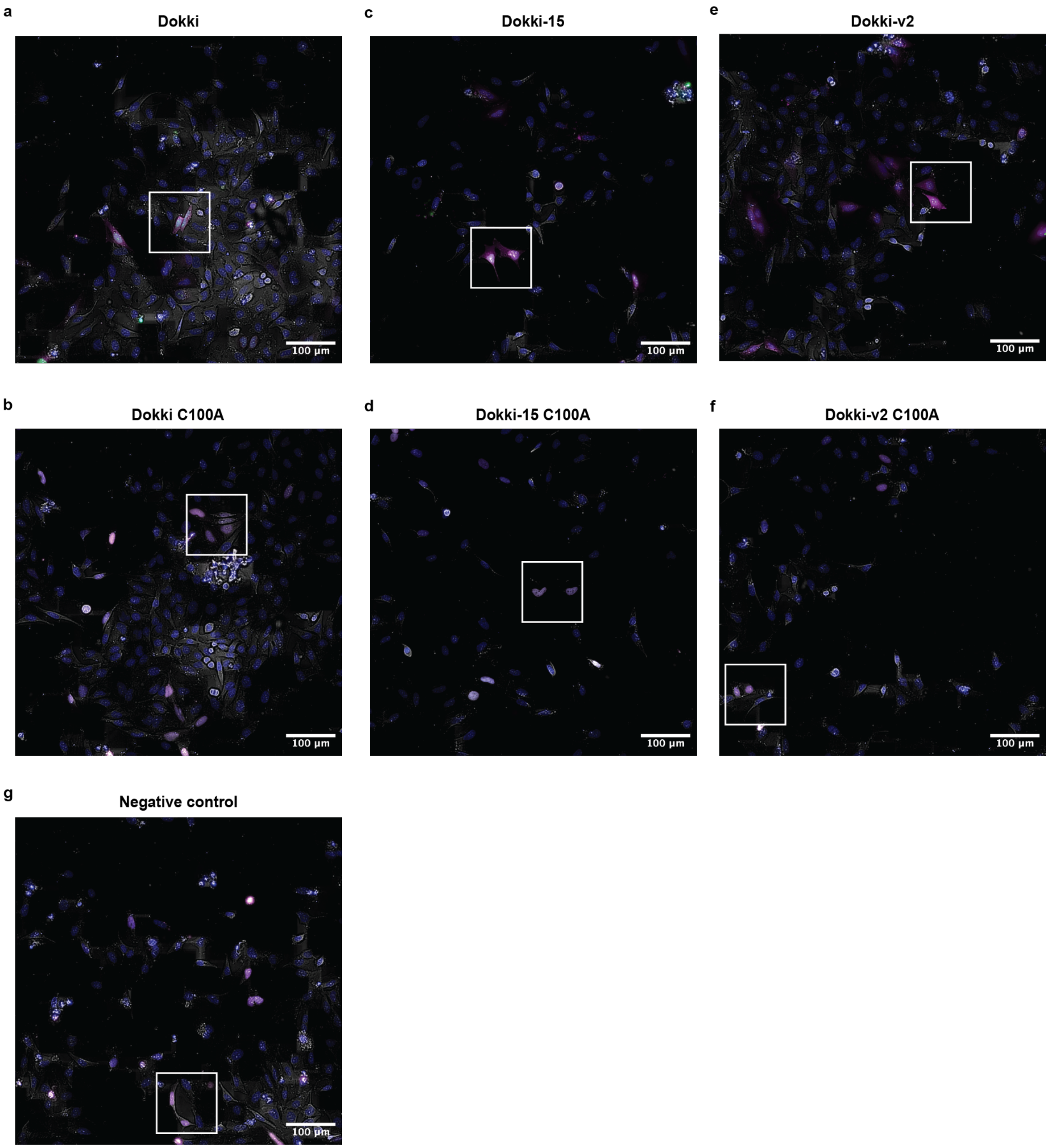

**Supplementary Fig. 3 | Raw image for cell-based reporter assay**

**a–g,** Representative fluorescence microscopy images of HeLa cells before crop. Cropped region is shown as white box **a,** Dokki; **b,** Dokk-15i; **c,** Dokki-v2; **d,** Dokki C100A; **e,** Dokki-15 C100A; **f,** Dokki-v2 C100A; **g,** Negative control.

**Supplementary information 1. pCool6 plasmid DNA sequence**

actcctgcattaggaagcagcccagtagtaggttgaggccgttgagcaccgccgccgcaaggaatggtgcatgcaaggagatggcgcccaacagtcccccggcca
cggggcctgccaccatacccacgccgaaacaagcgctcatgagcccgaagtggcgagcccgatcttccccatcggtgatgtcggcgatataggcgccagcaaccg
cacctgtggcgccggtgatgccggccacgatgcgtccggcgtagaggatcgagatcgatctcgatcccgcgaaattaatacgactcactataggggaattgtgagcg
gataacaattcccctctagaaataattttgtttaactttaagaaggagatatacatatgTCAGGAGGAGACCGAaagcgtattacagtgacagttgacagcg
acagctatcagttgctcaaggcatatatgatgtcaatatctccggtctggtaTgcacaaccaagaatgaagcccATGCAGTTTAAGGTTTACACCT
ATAAAAGAGAGAGCCGTTATCGTCTGTTTGTGGATGTACAGAGTGATATTATTGACACGCCCGGGCG
ACGGATGGTGATCCCCCTGGCCAGTGCACGTCTGCTGTCAGATAAAGTCTCCCGTGAACTTTACCCG
GTGGTGCATATCGGGGATGAAAGCTGGCGCATGATGACCACCGATATGGCCAGTGTGCCGGTtTCCG
TTATCGGGGAAGAAGTGGCTGATCTCAGCCACCGCGAAAATGACATCAAAAACGCCATTAACCTGA
TGTTCTGGGGAATAtgaACGGTCTCGTTCCGGCAGCGGCAGCGGCAGCATGGATAGCACCGAAGCGGT
GATTAAAGAATTTATGCGCTTTAAAGTGCATATGGAAGGCAGCATGAACGGCCATGAATTTGAAATT
GAAGGTGAAGGCGAAGGCCGCCCGTATGAAGGCACCCAGACCGCGAAACTGAAAGTGACCAAAGG
CGGCCCGCTGCCGTTTAGCTGGGATATTCTGAGCCCGCAGTTTATGTATGGCAGCCGCGCGTTTATTA
AACATCCGGCGGATATCCCGGATTATTGGAAACAGAGCTTTCCGGAAGGTTTTAAATGGGAACGCGT
GATGATCTTTGAAGATGGTGGCACCGTGAGCGTGACCCAGGATACCAGCCTGGAAGATGGCACCCT
GATTTATAAAGTGAAACTGCGCGGCGGCAACTTTCCGCCGGATGGCCCGGTGATGCAGAAACGCAC
CATGGGCTGGGAAGCGAGCACCGAACGCCTGTATCCGGAAGATGTGGTGCTGAAAGGCGATATTAA
AATGGCGCTGCGCCTGAAAGATGGCGGCCGCTATCTGGCGGATTTTAAAACCACCTATAAAGCGAA
GAAACCGGTGCAGATGCCGGGCGCCTTTAACATTGATCGCAAACTGGATATTACCAGCCATAACGA
AGATTATACCGTGGTGGAACAGTATGAACGCAGCGTGGCGCGCCATAGCACCGGCTCTcatcatcatcacca
TCATtaatgagatccggctgctaacaaagcccgaaaggaagctgagttggctgctgccaccgctgagcaataactagcataaccccttggggcctctaaacgggt
cttgaggggttttttgctgaaaggaggaactatatccggattggcgaatgggacgcgccctgtagcggcgcattaagcgcggcgggtgtggtggttacgcgcagcgt
gaccgctacacttgccagcgccctagcgcccgctcctttcgctttcttcccttcctttctcgccacgttcgccggctttccccgtcaagctctaaatcgggggctcccttta
gggttccgatttagtgctttacggcacctcgaccccaaaaaacttgattagggtgatggttcacgtagtgggccatcgccctgatagacggtttttcgccctttgacgttgg
agtccacgttctttaatagtggactcttgttccaaactggaacaacactcaaccctatctcggtctattcttttgatttataagggattttgccgatttcggcctattggttaaaa
aatgagctgatttaacaaaaatttaacgcgaattttaacaaaatattaacgtttacaatttcaggtggcacttttcggggaaatgtgcgcggaacccctatttgtttatttttcta
aatacattcaaatatgtatccgctcatgaattaattcttagaaaaactcatcgagcatcaaatgaaactgcaatttattcatatcaggattatcaataccatatttttgaaaaag
ccgtttctgtaatgaaggagaaaactcaccgaggcagttccataggatggcaagatcctggtatcggtctgcgattccgactcgtccaacatcaatacaacctattaattt
cccctcgtcaaaaataaggttatcaagtgagaaatcaccatgagtgacgactgaatccggtgagaatggcaaaagtttatgcatttctttccagacttgttcaacaggcca
gccattacgctcgtcatcaaaatcactcgcatcaaccaaaccgttattcattcgtgattgcgcctgagcgagacgaaatacgcgatcgctgttaaaaggacaattacaaa
caggaatcgaatgcaaccggcgcaggaacactgccagcgcatcaacaatattttcacctgaatcaggatattcttctaatacctggaatgctgttttcccggggatcgca
gtggtgagtaaccatgcatcatcaggagtacggataaaatgcttgatggtcggaagaggcataaattccgtcagccagtttagtctgaccatctcatctgtaacatcattg
gcaacgctacctttgccatgtttcagaaacaactctggcgcatcgggcttcccatacaatcgatagattgtcgcacctgattgcccgacattatcgcgagcccatttatac
ccatataaatcagcatccatgttggaatttaatcgcggcctagagcaagacgtttcccgttgaatatggctcataacaccccttgtattactgtttatgtaagcagacagtttt
attgttcatgaccaaaatcccttaacgtgagttttcgttccactgagcgtcagaccccgtagaaaagatcaaaggatcttcttgagatcctttttttctgcgcgtaatctgctg
cttgcaaacaaaaaaaccaccgctaccagcggtggtttgtttgccggatcaagagctaccaactctttttccgaaggtaactggcttcagcagagcgcagataccaaat
actgtccttctagtgtagccgtagttaggccaccacttcaagaactctgtagcaccgcctacatacctcgctctgctaatcctgttaccagtggctgctgccagtggcgat
aagtcgtgtcttaccgggttggactcaagacgatagttaccggataaggcgcagcggtcgggctgaacggggggttcgtgcacacagcccagcttggagcgaacg
acctacaccgaactgagatacctacagcgtgagctatgagaaagcgccacgcttcccgaagggagaaaggcggacaggtatccggtaagcggcagggtcggaac
aggagagcgcacgagggagcttccagggggaaacgcctggtatctttatagtcctgtcgggtttcgccacctctgacttgagcgtcgatttttgtgatgctcgtcaggg
gggcggagcctatggaaaaacgccagcaacgcggcctttttacggttcctggccttttgctggccttttgctcacatgttctttcctgcgttatcccctgattctgtggataa
ccgtattaccgcctttgagtgagctgataccgctcgccgcagccgaacgaccgagcgcagcgagtcagtgagcgaggaagcggaagagcgcctgatgcggtatttt
ctccttacgcatctgtgcggtatttcacaccgcatatatggtgcactctcagtacaatctgctctgatgccgcatagttaagccagtatacactccgctatcgctacgtgact
gggtcatggctgcgccccgacacccgccaacacccgctgacgcgccctgacgggcttgtctgctcccggcatccgcttacagacaagctgtgaccgtctccgggag
ctgcatgtgtcagaggttttcaccgtcatcaccgaaacgcgcgaggcagctgcggtaaagctcatcagcgtggtcgtgaagcgattcacagatgtctgcctgttcatcc

gcgtccagctcgttgagtttctccagaagcgttaatgtctggcttctgataaagcgggccatgttaagggcggtttttcctgtttggtcactgatgcctccgtgtaaggggg
atttctgttcatgggggtaatgataccgatgaaacgagagaggatgctcacgatacgggttactgatgatgaacatgcccggttactggaacgttgtgagggtaaaca
actggcggtatggatgcggcgggaccagagaaaaatcactcagggtcaatgccagcgcttcgttaatacagatgtaggtgttccacagggtagccagcagcatcctg
cgatgcagatccggaacataatggtgcagggcgctgacttccgcgtttccagactttacgaaacacggaaaccgaagaccattcatgttgttgctcaggtcgcagacg
ttttgcagcagcagtcgcttcacgttcgctcgcgtatcggtgattcattctgctaaccagtaaggcaaccccgccagcctagccgggtcctcaacgacaggagcacgat
catgcgcacccgtggggccgccatgccggcgataatggcctgcttctcgccgaaacgtttggtggcgggaccagtgacgaaggcttgagcgagggcgtgcaagat
tccgaataccgcaagcgacaggccgatcatcgtcgcgctccagcgaaagcggtcctcgccgaaaatgacccagagcgctgccggcacctgtcctacgagttgcat
gataaagaagacagtcataagtgcggcgacgatagtcatgccccgcgcccaccggaaggagctgactgggttgaaggctctcaagggcatcggtcgagatcccgg
tgcctaatgagtgagctaacttacattaattgcgttgcgctcactgcccgctttccagtcgggaaacctgtcgtgccagctgcattaatgaatcggccaacgcgcgggga
gaggcggtttgcgtattgggcgccagggtggtttttcttttcaccagtgagacgggcaacagctgattgcccttcaccgcctggccctgagagagttgcagcaagcggt
ccacgctggtttgccccagcaggcgaaaatcctgtttgatggtggttaacggcgggatataacatgagctgtcttcggtatcgtcgtatcccactaccgagatgtccgca
ccaacgcgcagcccggactcggtaatggcgcgcattgcgcccagcgccatctgatcgttggcaaccagcatcgcagtgggaacgatgccctcattcagcatttgcat
ggtttgttgaaaaccggacatggcactccagtcgccttcccgttccgctatcggctgaatttgattgcgagtgagatatttatgccagccagccagacgcagacgcgcc
gagacagaacttaatgggcccgctaacagcgcgatttgctggtgacccaatgcgaccagatgctccacgcccagtcgcgtaccgtcttcatgggagaaaataatact
gttgatgggtgtctggtcagagacatcaagaaataacgccggaacattagtgcaggcagcttccacagcaatggcatcctggtcatccagcggatagttaatgatcag
cccactgacgcgttgcgcgagaagattgtgcaccgccgctttacaggcttcgacgccgcttcgttctaccatcgacaccaccacgctggcacccagttgatcggcgc
gagatttaatcgccgcgacaatttgcgacggcgcgtgcagggccagactggaggtggcaacgccaatcagcaacgactgtttgcccgccagttgttgtgccacgcg
gttgggaatgtaattcagctccgccatcgccgcttccactttttcccgcgttttcgcagaaacgtggctggcctggttcaccacgcgggaaacggtctgataagagacac
cggcatactctgcgacatcgtataacgttactggtttcacattcaccaccctgaattgactctcttccgggcgctatcatgccataccgcgaaaggttttgcgccattcgat
ggtgtccgggatctcgacgctctcccttatgcg

**Supplementary information 2. ZJ101 plasmid DNA sequence**

gaagatcctttgatcttttctacggggtctgacgctcagtggaacgaaaactcacgttaagggattttggtcatgagattatcaaaaaggatcttcacctagatccttttaaat
taaaaatgaagttttaaatcaatctaaagtatatatgagtaaacttggtctgacagttaccaatgcttaatcagtgaggcacctatctcagcgatctgtctatttcgttcatcca
tagttgcctgactccccgtcgtgtagataactacgatacgggagggcttaccatctggccccagtgctgcaatgataccgcgGgacccacgctcaccggctccagatt
tatcagcaataaaccagccagccggaagggccgagcgcagaagtggtcctgcaactttatccgcctccatccagtctattaattgttgccgggaagctagagtaagta
gttcgccagttaatagtttgcgcaacgttgttgccattgctacaggcatcgtggtgtcacgctcgtcgtttggtatggcttcattcagctccggttcccaacgatcaaggcg
agttacatgatcccccatgttgtgcaaaaaagcggttagctccttcggtcctccgatcgttgtcagaagtaagttggccgcagtgttatcactcatggttatggcagcactg
cataattctcttactgtcatgccatccgtaagatgcttttctgtgactggtgagtactcaaccaagtcattctgagaatagtgtatgcggcgaccgagttgctcttgcccggc
gtcaatacgggataataccgcgccacatagcagaactttaaaagtgctcatcattggaaaacgttcttcggggcgaaaactctcaaggatcttaccgctgttgagatcca
gttcgatgtaacccactcgtgcacccaactgatcttcagcatcttttactttcaccagcgtttctgggtgagcaaaaacaggaaggcaaaatgccgcaaaaaagggaata
agggcgacacggaaatgttgaatactcatactcttcctttttcaatattattgaagcatttatcagggttattgtctcatgagcggatacatatttgaatgtatttagaaaaataa
acaaataggggttccgcgcacatttccccgaaaagtgccacctgacgtggatcccctgagggggcccccatgggctagaggatccggcctcggcctctgcataaat
aaaaaaaattagtcagccatgagcttggcccattgcatacgttgtatccatatcataatatgtacatttatattggctcatgtccaacattaccgccatgttgacattgattatt
gactagttattaatagtaatcaattacggggtcattagttcatagcccatatatggagttccgcgttacataacttacggtaaatggcccgcctggctgaccgcccaacga
cccccgcccattgacgtcaataatgacgtatgttcccatagtaacgccaatagggactttccattgacgtcaatgggtggagtatttacggtaaactgcccacttggcagt
acatcaagtgtatcatatgccaagtacgccccctattgacgtcaatgacggtaaatggcccgcctggcattatgcccagtacatgaccttatgggactttcctacttggca
gtacatctacgtattagtcatcgctattaccatggtgatgcggttttggcagtacatcaatgggcgtggatagcggtttgactcacggggatttccaagtctccaccccatt
gacgtcaatgggagtttgttttggcaccaaaatcaacgggactttccaaaatgtcgtaacaactccgccccattgacgcaaatgggcggtaggcgtgtacggtgggag
gtctatataagcagagctcgtttagtgaaccgtcagatcgcctggagacgccatccacgctgttttgacctccatagaagacaccgggaccgatccagcctcccctcga
agcttacatgtggtaccgagctcggatcctgagaacttcagggtgagtctatgggacccttgatgttttctttccccttcttttctatggttaagttcatgtcataggaagggg
agaagtaacagggtacacatattgaccaaatcagggtaattttgcatttgtaattttaaaaaatgctttcttcttttaatatacttttttgtttatcttatttctaatactttccctaatct
ctttctttcagggcaataatgatacaatgtatcatgcctctttgcaccattctaaagaataacagtgataatttctgggttaaggcaatagcaatatttctgcatataaatatttct
gcatataaattgtaactgatgtaagaggtttcatattgctaatagcagctacaatccagctaccattctgcttttattttatggttgggataaggctggattattctgagtccaa
gctaggcccttttgctaatcatgttcatacctcttatcttcctcccacagctcctgggcaacgtgctggtctgtgtgctggcccatcactttggcaaagcacgtgagatctga
attctgacactAAGGTTGAGACCgccaccaagcgtattacagtgacagttgacagcgacagctatcagttgctcaaggcatatatgatgtcaatatctccggt

ctggtaTgcacaaccaagaatgaagcccATGCAGTTTAAGGTTTACACCTATAAAAGAGAGAGCCGTTATCGTCTGT
TTGTGGATGTACAGAGTGATATTATTGACACGCCCGGGCGACGGATGGTGATCCCCCTGGCCAGTGC
ACGTCTGCTGTCAGATAAAGTCTCCCGTGAACTTTACCCGGTGGTGCATATCGGGGATGAAAGCTGG
CGCATGATGACCACCGATATGGCCAGTGTGCCGGTtTCCGTTATCGGGGAAGAAGTGGCTGATCTCA
GCCACCGCGAAAATGACATCAAAAACGCCATTAACCTGATGTTCTGGGGAATAtgaGGTCTCcCCTActc
aaatcctgcacaacagattcttcatgtttggaccaaatcaacttgtgataccatgctcaaagaggcctcaattatatttgagtttttaatttttatgaaaaaaaaaaaaaaaaa
cggaattcaccccaccagtgcaggctgcctatcagaaagtggtggctggtgtggctaatgccctggcccacaagtatcactaagctcgctttcttgctgtccaatttctat
taaaggttcctttgttccctaagtccaactactaaactgggggatattatgaagggccttgagcatctggattctgcctaataaaaaacatttattttcattgcaatgatgtattt
aaattatttctgaatattttactaaaaagggaatgtgggaggtcagtgcatttaaaacataaagaaatgaagagctagttcaaaccttgggaaaatacactatatcttaaact
ccatgaaagaaggtgaggctgcaaacagctaatgcacattggcaacagcccctgatgcctatgccttattcatccctcagaaaaggattcaagtagaggcttgatttgg
aggttaaagttttgctatgctgtattttacattacttattgttttagctgtcctcatgaatgtcttttcactacccatttgcttatcctgcatctctcagccttgactccactcagttctc
ttgcttagagataccacctttcccctgaagtgttccttccatgttttacggcgagatggtttctcctcgcctggccactcagccttagttgtctctgttgtcttatagaggtctac
ttgaagaaggaaaaacagggggcatggtttgactgtcctgtgagcccttcttccctgcctcccccactcacagtgacccggaatccctcgacatggcagtctagcacta
gtgcggccgcagatctgcttcctcgctcactgactcgctgcgctcggtcgttcggctgcggcgagcggtatcagctcactcaaaggcggtaatacggttatccacaga
atcaggggataacgcaggaaagaacatgtgagcaaaaggccagcaaaaggccaggaaccgtaaaaaggccgcgttgctggcgtttttccataggctccgcccccc
tgacgagcatcacaaaaatcgacgctcaagtcagaggtggcgaaacccgacaggactataaagataccaggcgtttccccctggaagctccctcgtgcgctctcctg
ttccgaccctgccgcttaccggatacctgtccgcctttctcccttcgggaagcgtggcgctttctcatagctcacgctgtaggtatctcagttcggtgtaggtcgttcgctc
caagctgggctgtgtgcacgaaccccccgttcagcccgaccgctgcgccttatccggtaactatcgtcttgagtccaacccggtaagacacgacttatcgccactggc
agcagccactggtaacaggattagcagagcgaggtatgtaggcggtgctacagagttcttgaagtggtggcctaactacggctacactagaagaacagtatttggtat
ctgcgctctgctgaagccagttaccttcggaaaaagagttggtagctcttgatccggcaaacaaaccaccgctggtagcggtggtttttttgtttgcaagcagcagattac
gcgcagaaaaaaaggatctcaa
